# Gut microbiome shifts in chronic systolic heart failure are associated with disease severity and clinical improvement

**DOI:** 10.1101/2024.08.06.606872

**Authors:** Petra Mamic, Handuo Shi, Wenyu Zhou, Nasim Bararpour, Kevin Contrepois, Heyjun Park, Monika Avina, Sophia Miryam Schüssler-Fiorenza Rose, Paul A. Heidenreich, Kiran Kaur Khush, Michael B. Fowler, W. H. Wilson Tang, Karim Sallam, Justin Sonnenburg, Kerwyn Casey Huang, Michael P. Snyder

**Author notes:** These authors contributed equally (Petra Mamic, Handuo Shi, Kerwyn Casey Huang, Michael P. Snyder).

## Abstract

Chronic systolic heart failure (HF) is a prevalent and morbid disease with marked variability in its progression and response to therapies. The gut microbiome may play a role in pathophysiology and progression of chronic HF, but clinical studies investigating relationships between the two are lacking. We analyzed the gut microbiome in a cohort of adults with chronic systolic HF caused by non-ischemic cardiomyopathy (*n*=59) using multi-omics profiling and, in some cases, longitudinal sampling. We identified microbiome differences compared to healthy subjects (*n*=50) and associated these differences with host metabolites, inflammatory markers and physiology. We found depletion of the anti-inflammatory probiotic *Bifidobacterium* and the associated short chain fatty acid producing and formaldehyde detoxifying pathways in the chronic HF cohort. We also discovered HF-specific microbiome-host immunome interactions. In addition to identifying several taxa and microbial pathways broadly associated with HF disease severity, we found significant links between *Bifidobacterium* and clinical HF improvement over time. Gut microbiome-host multi-omic data integration revealed a close association between *Bifidobacterium* and circulating metabolites previously implicated in cardiovascular physiology (e.g., malonic acid), thus pointing to potential mechanisms through which *Bifidobacterium* may affect chronic HF physiology. Our results suggest that *Bifidobacterium* may serve as a biomarker for chronic HF trajectory as well as suggest potential novel therapeutic interventions strategies.

## INTRODUCTION

Over 6 million adults in the U.S. and 60 million adults worldwide live with chronic heart failure (HF)^1^. Despite advances in management, chronic HF is still associated with significant morbidity and mortality^2^. Additionally, HF disease course, even with appropriate therapy, is variable^3^, and our ability to predict it on individual level is limited, partly due to incomplete understanding of disease mechanisms and factors that contribute to disease progression.

Recent studies suggest that the gut microbiome, a diverse community of microbes with myriad links to human health^4^, plays an important role in HF pathophysiology. Chronic HF has been associated with gut microbiome disturbances^5–11^, including expansion of potential gut pathogens^7^ at the expense of anti-inflammatory microbial taxa^11–13^. Heart failure-associated alterations to gut microbiome composition are accompanied by functional shifts, including depletion of pathways for biosynthesis of short chain fatty acids (SCFAs)^5,6^ and enrichment of lipopolysaccharide biosynthetic pathways^5^. In animal models these microbial products affect various aspects of HF physiology; for example, SCFAs – byproducts of microbial fermentation of dietary fiber – regulate cardiac remodeling, vascular tone, and regulatory T-cell proliferation^14–18^, in addition to serving as fuel for the failing heart^19^. However, mechanistic links between the gut microbiome and HF pathophysiology in humans remain unclear, with previous studies limited by small sample size, cross-sectional design, and lack of functional data such as metabolomics.

Here, we report the first gut microbiome-focused, longitudinal, multi-omics study in patients with chronic systolic HF related to non-ischemic cardiomyopathy (NICM). We identify HF-associated taxonomic and functional gut microbiome signatures, and gut microbiome, circulating metabolome, and immunome signatures of HF severity and clinical improvement over time. These findings reveal potential mechanisms through which gut microbes may affect host physiology in chronic systolic HF.

## RESULTS

### Cohort description

Figure 1 highlights the study design, data collected, and analyses performed. Briefly, we enrolled a cohort of 59 adults with chronic HF and 50 healthy controls. Table 1 outlines relevant demographics, comorbidities, medications, and clinical laboratory values at baseline for the two cohorts. The mean age of study participants was 53.7 years, mean body mass index (BMI) was 27.9 kg/m^2^, and approximately two thirds of each cohort self-identified as white. Women were significantly underrepresented in the HF relative to the healthy cohort (22% versus 52%, *p*=0.001).

**Figure 1:**
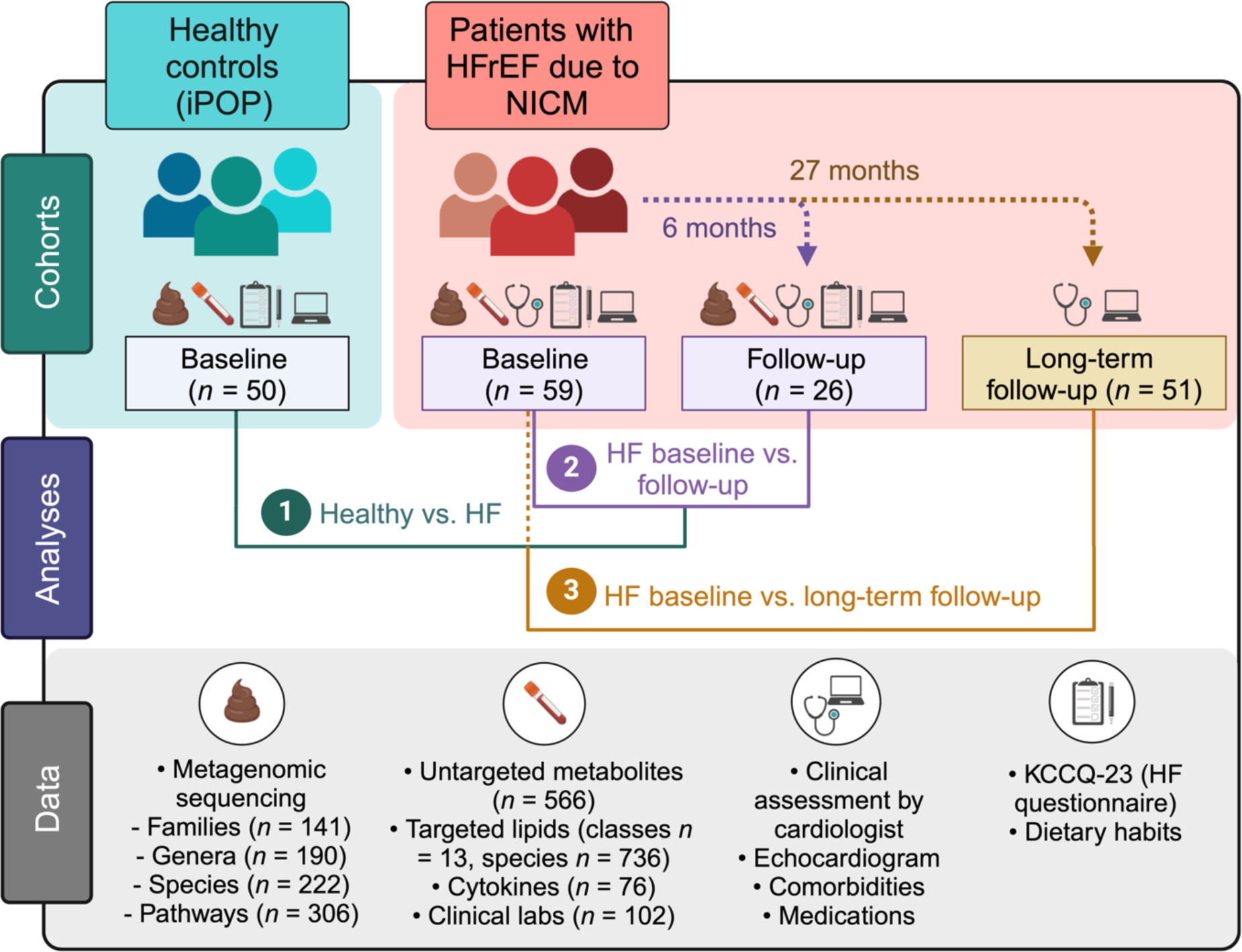
Design of multi-omic study with longitudinal profiling and clinical phenotyping of a cohort of 59 adults with chronic heart failure with reduced ejection fraction due to non-ischemic cardiomyopathy. Adults with stable chronic heart failure with reduced ejection fraction (HFrEF) due to non-ischemic cardiomyopathy (NICM) (*n*=59) were recruited for the study. At baseline visit each participant underwent multi-omic profiling, including (a) whole genome metagenomic sequencing of microbial DNA extracted from stool, (b) untargeted metabolomics and (c) targeted lipidomics from host plasma, (d) 76-cytokine Luminex immune panel from host serum, and (e) extensive clinical lab profiling. All individuals with chronic heart failure (HF) were additionally assessed by their regular HF cardiologist within days of their baseline study visit. Clinical data (patient functional cardiac status and symptomatology, echocardiographic parameters, comorbidities, and medications) was extracted from the electronic medical record (EMR). Study participants additionally completed questionnaires related to their cardiac functional status and HF symptoms (Kansas City Cardiomyopathy Questionnaire-23 (KCCQ-23)) and habitual diet. A follow-up visit was completed by 26 participants after 6 months on average after their baseline visit, at which point they underwent repeat multi-omic profiling, clinical assessment, and completed questionnaires. For 51 of the patients with at least a baseline visit, limited clinical data (clinical status and limited echocardiographic data) was available and extracted from the EMR. Healthy subjects (*n*=50) were previously recruited for the integrated Personal Omics Profiling (iPOP) study (one of the three core studies of the National Institutes of Health Integrative Human Microbiome Project^32,76^), during which they were profiled extensively on a quarterly basis. Stool and serum samples, and questionnaire data from the most recent healthy visit were included for this study. (Figure created with BioRender.com.)

**Table 1:**
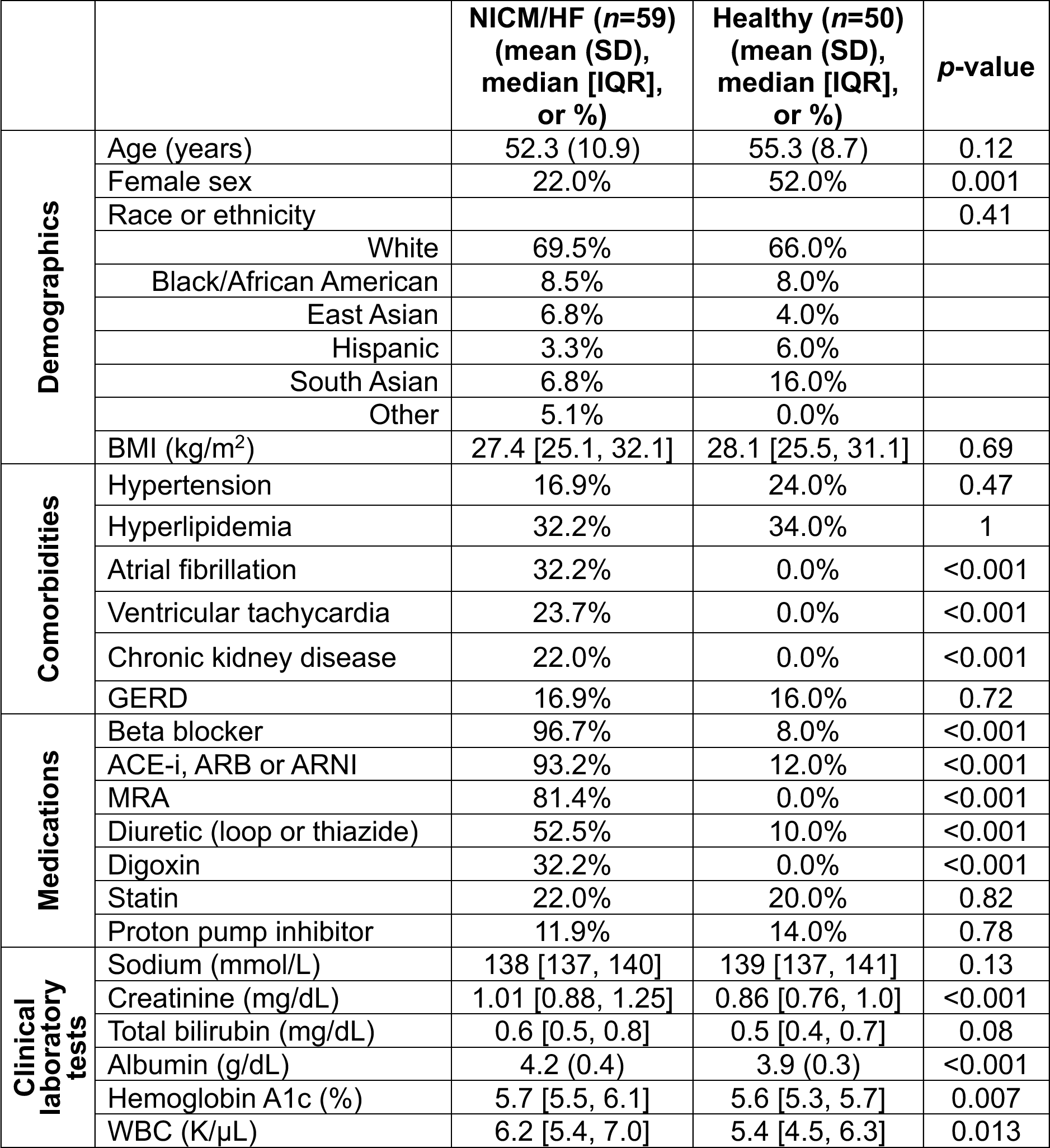
Baseline characteristics of individuals from the NICM/HF and healthy cohorts. Abbreviations: NICM: non-ischemic cardiomyopathy; HF: heart failure; SD: standard deviation; IQR: interquartile range; BMI: body mass index; GERD: gastroesophageal reflux disease; ACE-i: angiotensin-converting enzyme inhibitor; ARB: angiotensin II receptor blocker; ARNI: angiotensin receptor-neprilysin inhibitor; WBC: white blood cell count. Student’s t-tests were used for between-group comparison of normally distributed continuous variables, Mann-Whitney U tests were used for continuous variables with skewed distribution, and Fisher’s exact tests were used for categorical variables.

|  |  | NICM/HF (*n*=59)<br>(mean (SD),<br>median [IQR],<br>or %) | Healthy (*n*=50)<br>(mean (SD),<br>median [IQR],<br>or %) | *p*-value |
| --- | --- | --- | --- | --- |
| Demographics | Age (years) | 52.3 (10.9) | 55.3 (8.7) | 0.12 |
|  | Female sex | 22.0% | 52.0% | 0.001 |
|  | Race or ethnicity |  |  | 0.41 |
|  | White | 69.5% | 66.0% |  |
|  | Black/African American | 8.5% | 8.0% |  |
|  | East Asian | 6.8% | 4.0% |  |
|  | Hispanic | 3.3% | 6.0% |  |
|  | South Asian | 6.8% | 16.0% |  |
|  | Other | 5.1% | 0.0% |  |
|  | BMI (kg/m<sup>2</sup>) | 27.4 [25.1, 32.1] | 28.1 [25.5, 31.1] | 0.69 |
| Comorbidities | Hypertension | 16.9% | 24.0% | 0.47 |
|  | Hyperlipidemia | 32.2% | 34.0% | 1 |
|  | Atrial fibrillation | 32.2% | 0.0% | <0.001 |
|  | Ventricular tachycardia | 23.7% | 0.0% | <0.001 |
|  | Chronic kidney disease | 22.0% | 0.0% | <0.001 |
|  | GERD | 16.9% | 16.0% | 0.72 |
| Medications | Beta blocker | 96.7% | 8.0% | <0.001 |
|  | ACE-i, ARB or ARNI | 93.2% | 12.0% | <0.001 |
|  | MRA | 81.4% | 0.0% | <0.001 |
|  | Diuretic (loop or thiazide) | 52.5% | 10.0% | <0.001 |
|  | Digoxin | 32.2% | 0.0% | <0.001 |
|  | Statin | 22.0% | 20.0% | 0.82 |
|  | Proton pump inhibitor | 11.9% | 14.0% | 0.78 |
| Clinical laboratory tests | Sodium (mmol/L) | 138 [137, 140] | 139 [137, 141] | 0.13 |
|  | Creatinine (mg/dL) | 1.01 [0.88, 1.25] | 0.86 [0.76, 1.0] | <0.001 |
|  | Total bilirubin (mg/dL) | 0.6 [0.5, 0.8] | 0.5 [0.4, 0.7] | 0.08 |
|  | Albumin (g/dL) | 4.2 (0.4) | 3.9 (0.3) | <0.001 |
|  | Hemoglobin A1c (%) | 5.7 [5.5, 6.1] | 5.6 [5.3, 5.7] | 0.007 |
| | WBC (K/μL) | 6.2 [5.4, 7.0] | 5.4 [4.5, 6.3] | 0.013 |

Table 2 summarizes relevant baseline cardiomyopathy and HF phenotype data for the HF cohort. In this cohort of NICM patients, 44.1% had idiopathic and 35.6% had familial cardiomyopathy. The mean left ventricular ejection fraction (LVEF) was markedly reduced, which was associated with a mildly to moderately dilated left ventricle (LV).

**Table 2:**
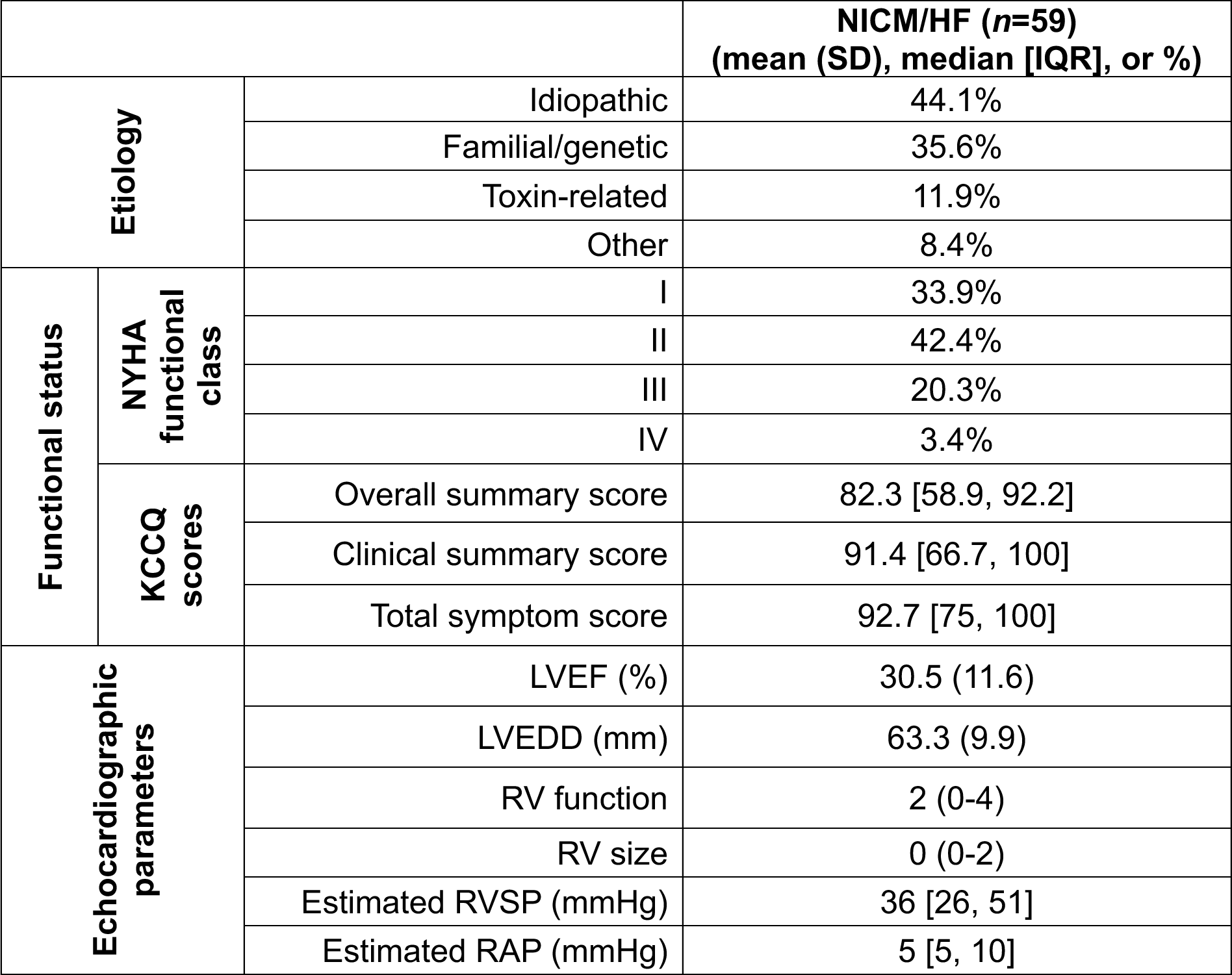
Cardiomyopathy and heart failure phenotype for study participants at baseline. Abbreviations: NICM: non-ischemic cardiomyopathy; HF: heart failure; SD: standard deviation; IQR: interquartile range; NYHA: New York Heart Association; KCCQ: Kansas City Cardiomyopathy Questionnaire; LVEF: left ventricular ejection fraction; LVEDD: left ventricular end-diastolic dimension; RV: right ventricular; RVSP: right ventricular systolic pressure; RAP: right atrial pressure. Student’s t-tests were used for between-group comparison of normally distributed continuous variables, Mann-Whitney U tests were used for continuous variables with a skewed distribution, and Fisher’s exact tests were used for categorical variables.

Right ventricular (RV) dysfunction on echocardiograms was present in >50% of the cohort. Most of the individuals with HF were clinically well compensated, with New York Heart Association (NYHA) functional class I and II symptoms, and a relatively high median Kansas City Cardiomyopathy Questionnaire-23 (KCCQ-23) scores.

Compared to the healthy cohort, patients with chronic HF had more comorbidities and were prescribed more medications (Table 1); as expected, the most notable differences were in cardiac medications, which comprise guideline-directed medical therapy for individuals with systolic HF. Table S1 lists complete between-group differences in comorbidities, medications, and labs.

Twenty-six patients from the HF cohort returned for a follow-up visit at a mean of 6 months after their baseline visit (one patient returned for two follow-up visits at 3 and 7 months). Clinical follow-up data were extracted from the electronic medical record (EMR) and were available for 51 patients at an average of 27 months following their baseline visit. During study follow-up, 12 patients experienced a poor clinical outcome (a composite of being listed for or receiving a heart transplant, implantation of a durable left ventricular assist device (LVAD), transition to hospice or death). Fourteen individuals experienced clinical improvement, measured by improvement in NYHA class, LVEF or KCCQ-23. The longitudinal nature of this outpatient systolic HF cohort, focused on individuals with NICM, combined with our ability to capture data on individual clinical trajectories and to integrate this information with comprehensive gut microbiome and host multi-omics data provides a unique resource for mechanistic discovery.

### Gut microbiome taxonomic composition differs significantly in patients with chronic HF compared with healthy subjects

To profile gut microbiome composition across our cohorts, we performed metagenomic sequencing of stool samples. Of note, HF and healthy cohorts collected stool samples using two different methods, but we did not find a significant batch effect between them (Methods; Figure S1; Table S2). We fit data with a linear mixed-effects model that accounted for fixed effects due to subject age, sex, race, BMI, and reported dietary patterns, as well as a random effect for subject (Methods). Compared to healthy subjects, microbiome alpha diversity was significantly lower in patients with chronic HF (Fig. S2a). Beta diversity analysis revealed global gut microbiome differences between the two cohorts (*p*=0.001, permutational multivariate analysis of variance (PERMANOVA) test; Fig. S2b).

We next sought to identify differentially abundant taxa between the two cohorts (Table S3). The *Lachnospiraceae* and *Bifidobacteriaceae* families and the genera *Bifidobacterium, Anaerobutyricum, Anaerostipes, Lachnospira,* and *Blautia* (all Benjamini-Hochberg-adjusted *p*-value (*p*_adj_)<0.001) as well as *Longibaculum* (*p*_adj_=0.003) were depleted in patients with chronic HF (Fig. 2a,b). Many of these taxa exert beneficial anti-inflammatory effects on the host through SCFA production, largely from available precursors such as dietary fiber^20^. By contrast, *Sutterella* (*p*_adj_=0.064) (Fig. 2b), *Escherichia* (*p*_adj_=0.107), and *Prevotella* (*p*_adj_=0.114; Fig. S2c) were enriched in the HF cohort. Members of these genera can have pro-inflammatory effects^21,22^, and many *Escherichia* species are pathogenic facultative anaerobes. Most of the >80 differentially abundant species were depleted in the HF cohort (Table S3), including all four of the *Bifidobacterium* species detected in our samples. Ten of 11 *Prevotella* species and prevalent gut commensals such as *Bacteroides thetaiotaomicron* (*p*_adj_=0.057; Fig. S2c) and *Escherichia coli* (*p*_adj_=0.062) were enriched in the HF cohort. In sum, our taxonomic analysis indicates that the gut microbiome of patients with chronic HF differs significantly from that of healthy subjects and is characterized by profound loss of anti-inflammatory bacteria and enrichment of microbes associated with flaring inflammation both within the gut and systemically.

**Figure 2:**
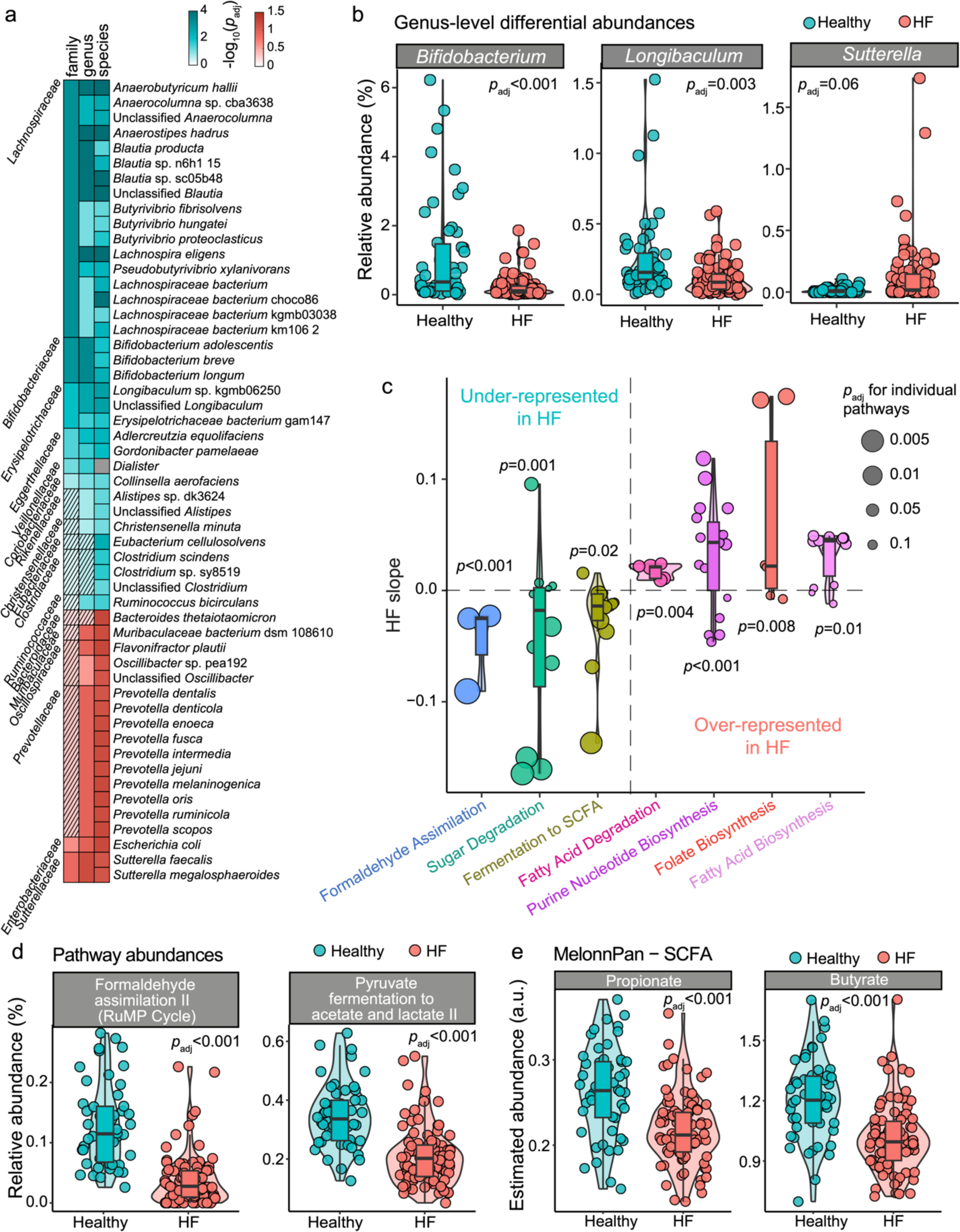
Chronic heart failure is associated with distinct taxonomic and functional gut microbiome shifts. **a**, Differentially abundant taxa between patients with chronic heart failure (HF) (*n*=59) compared to healthy subjects (*n*=50), with representative genera (*Bifidobacterium* and *Longibaculum* depleted, *Sutterella* enriched in HF) shown in **b**; **c**, Gene set enrichment analysis (GSEA) highlighted gut microbial pathways that are systematically over- or under-represented in the HF-associated gut microbiome, with key pathways shown in **d**; **e**, Consistent with the GSEA results, MelonnPan predicted significantly decreased gut microbiome-wide output of short chain fatty acids (SCFAs) butyrate and propionate. **a**-**b**, **d**-**e**, Individual microbiome features were modeled as dependent continuous variables in a linear mixed model with age, sex, race, body mass index (BMI), dietary patterns and categorical HF as fixed effects, and subject as random effect. **c**, For GSEA, *p*-value of each pathway group was estimated via 1,000 permutations. To adjust for multiple hypothesis testing, Benjamini-Hochberg method was used throughout (**a**-**e**). Abbreviations: HF = heart failure; SCFA = short chain fatty acid.

### Functional analyses indicate decreased SCFA biosynthetic potential in chronic HF

Based on the taxonomic shifts, we hypothesized that chronic HF-associated gut microbiome would show significant alterations in microbial metabolic pathways. Indeed, the abundance of gut microbial pathways differed significantly between the two cohorts (*p*=0.001, PERMANOVA test, Methods; Fig. S2d). Of the 306 annotated metabolic pathways, 100 were differentially abundant (Table S3). Several SCFA biosynthetic pathways were depleted in patients with chronic HF (Fig. 2c,d). Interestingly, some of the most depleted pathways were involved in formaldehyde detoxification (RUMP-PWY, PWY-1861, P185-PWY; all *p*_adj_<0.001; Fig. 2c,d, S2e). The pathway for methanogenesis from SCFA acetate (Fig. S2e) and pathways for L-arginine biosynthesis (all *p*_adj_<0.005) were also depleted in the HF cohort. Chronic HF cohort was enriched in microbial pathways related to biosynthesis of lipopolysaccharide (LPS) (PWY-1269 (*p*_adj_=0.002; Fig. S2e) and DTPRHAMSYN-PWY (*p*_adj_=0.074)), an integral component of the outer membrane of the Gram-negative microorganisms that incites the host inflammatory response^23–25^.

To systematically identify functional groups over- or under-represented in the HF cohort, we performed gene set enrichment analysis (GSEA^26^, Methods) on the microbiome metabolic pathways. Consistent with our individual observations, we found that fermentative and formaldehyde detoxification pathways were systematically depleted, while lipid metabolism, folate, and purine nucleotide biosynthetic pathways were enriched in patients with chronic HF (Fig. 2c, Table S3). This result suggests an overall shift from anaerobic carbohydrate to oxidative lipid metabolism, which has been associated with several disease states^18,27,28^. To predict microbially produced metabolites in the gut, we next applied the MelonnPan pipeline^29^ (Methods) to our metagenomics data. Consistent with our GSEA results, the predicted levels of several SCFAs (e.g., butyrate and propionate) were significantly lower in HF patients (Fig. 2e; Table S3). Altogether, our results suggest profound gut microbiome functional shifts in the context of chronic HF that may have systemic host effects.

### Chronic HF alters patterns of gut microbiome-host immune system interactions

Chronic HF has been associated with low-grade systemic inflammation that contributes to disease pathophysiology, progression, and complications^30^. The gut microbiome regulates local and systemic immunity through close host-microbiome interactions^31^. Given these connections, we investigated chronic HF-associated cytokine signatures and their associations with the gut microbiome. There were no significant differences in overall circulating immune profiles between the HF and healthy cohort (*p*=0.06, PERMANOVA; Table S4). However, our multivariate linear mixed model identified differences in individual cytokines with more apparent differences in patients with more severe HF (NYHA class III or IV, *n*=14) compared with healthy subjects (Table S4, Fig. S3a). Relatively modest differences are likely attributable to our HF cohort being overall well-compensated clinically and a higher prevalence of insulin resistance within the healthy cohort^32^.

We next investigated whether some of the gut microbiome-cytokine associations were cohort specific. Compared to the HF cohort, healthy subjects had more significant microbiome-cytokine correlations (Fig. S3b). Correlation patterns differed between the two groups, both for microbial pathways (Fig. 3a) and genera (Fig. S3c). One of the microbiome pathway clusters that showed a cohort-specific association with cytokines comprised L-arginine and ornithine biosynthetic pathways (Fig. 3a, marked with ***). These closely related amino acids affect gut permeability and host inflammatory state^33^.

**Figure 3:**
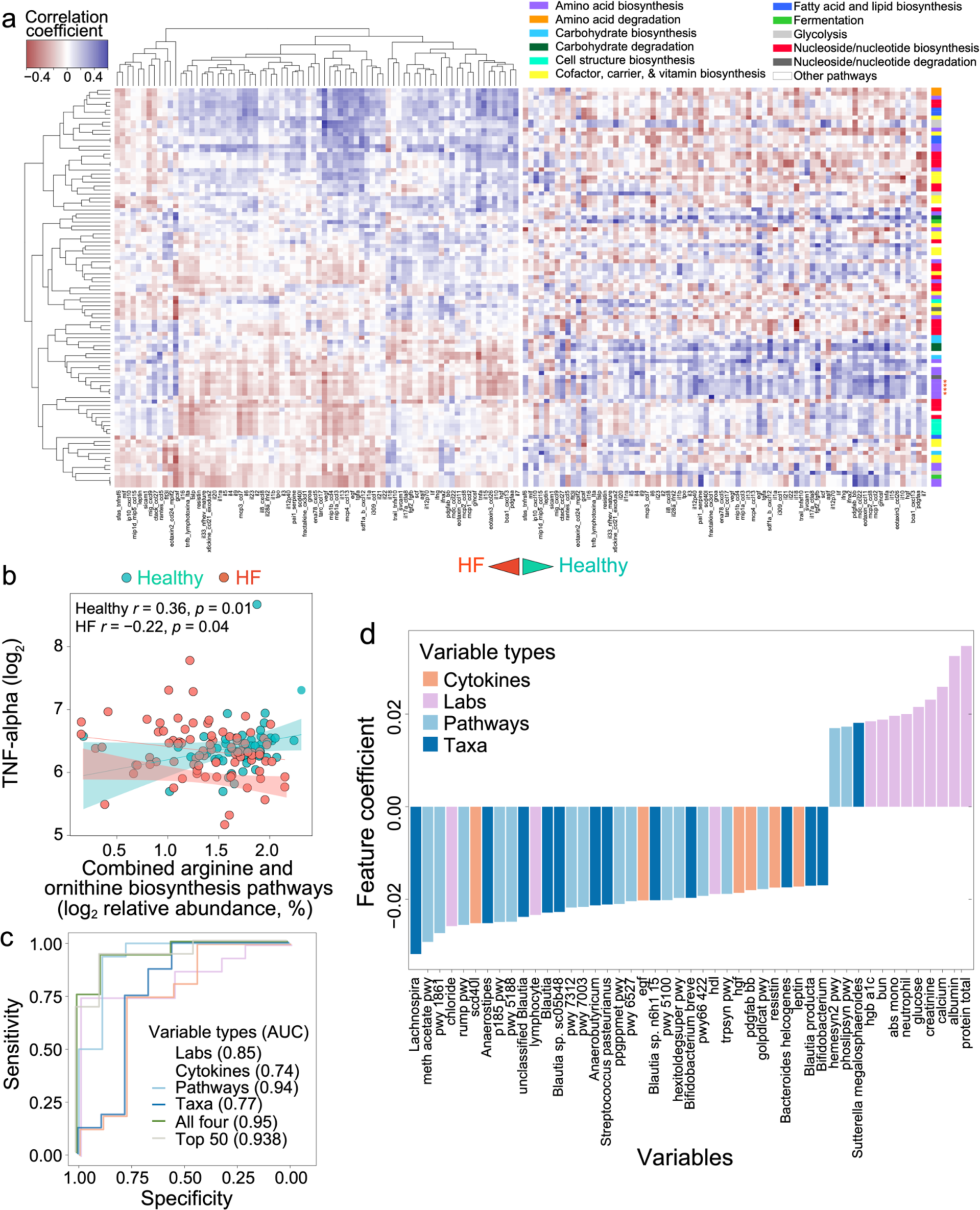
Gut microbiome-host cytokine connections are cohort-specific. **a**, Heatmap for correlations between top 100 most abundant microbial pathways (rows) and cytokines (*n*=76; columns). Pairwise Spearman correlations were calculated first, after which hierarchical clustering was performed (both on microbiome and cytokine levels) for the samples from the HF cohort (*n*=58) (left panel). The hierarchical cluster structure was then applied to the pairwise Spearman correlations between microbiome pathways and cytokines within the healthy cohort (*n*=48) (right panel). **b**, One of the microbiome pathway clusters that showed the most divergent, cohort-specific correlations with cytokines was the cluster of five pathways for arginine and ornithine biosynthesis (PWY-7400, ARGSYN-PWY, ARGSYNBSUB-PWY, PWY-5154, and GLUTORN-PWY) (marked with *** in **a**). The relative abundance of these pathways was aggregated, with panel **b** showing its cohort-specific association with TNF-⍺, the chief pro-inflammatory cytokine (Spearman correlation). Aggregated pathways were log_2_-transformed for visualization purposes. **c**, Machine learning algorithm was employed to investigate which data type was best at HF versus healthy classification. The algorithm included clinical laboratory data (*n*=53), cytokines (*n*=76), microbiome taxonomic (all ranks; *n*=553) and functional pathway data (*n*=306), from all visits (*n*=133). Variables were first standardized and 80% of the entire dataset were randomly selected from stratified groups as the training set with the rest as the holdout set. Lasso and elastic-net models were fitted for regression on standardized multi-omic variables using coordinate descent as implemented in R ‘glmnet’ and ‘caret’ packages. Panel **c** shows that microbiome pathways had by far the best classification performance (AUC 0.94), which was nearly equivalent to the combined performance of all four data types (AUC 0.95), and equivalent to the performance of the top 50 individual features (AUC 0.94). Top 50 individual features, of which more than half are microbiome related and include genus *Bifidobacterium*, are shown in **d**.

They are also precursors for biosynthesis of nitric oxide and polyamines, respectively, both of which have been associated with HF pathophysiology^34,35^. Notably, several of these pathways were depleted in our HF cohort (Table S3). It is not known to what degree gut microbiome arginine production contributes to the host arginine pool, but diminished global arginine bioavailability has been associated with more severe HF^36^. Arginine/ornithine biosynthetic potential was negatively correlated with the circulating levels of TNF-⍺ in our HF cohort, with the opposite correlation seen among healthy subjects (Fig. 3b). Although the magnitude of these correlations is low (possibly due to gut microbiome variability and cohort size), these findings suggest that gut microbiome-host immune system interactions are altered in chronic HF and may affect relevant disease processes.

### Gut microbial pathways are excellent classifiers of chronic HF

Since different types of molecules can reflect distinct facets of disease, we sought to understand how well our multi-omic datasets distinguished chronic HF and healthy states. We used clinical laboratory data (*n*=53), cytokines (*n*=76), and microbiome taxa (all ranks; *n*=553) and pathway data (*n*=306) to develop chronic HF classification models through machine learning (Methods). The entire multi-omic model exhibited excellent classification accuracy (area under the curve (AUC=0.95) (Fig. 3c). Notably, similar accuracy was achieved with only the microbiome pathway dataset or when only the top 50 most significant classification features were included (both AUC=0.94) (Fig. 3c). Of these features, 32 represented gut microbiome taxa (e.g., genera *Bifidobacterium*, *Lachnospira*, *Anaerostipes*, and *Blautia*) or functions (e.g., methanogenesis from SCFA acetate and formaldehyde detoxification) (Fig. 3d). These findings further support that a small number of multi-omic features – mostly gut microbiome-related – may be sufficient to classify chronic HF.

### Individual gut microbial taxa and pathways are associated with chronic HF severity and poor clinical outcome

Given the previously suggested involvement of gut microbial metabolic products (e.g., SCFAs) in HF pathophysiology^14–19^, we sought to determine if individual microbial taxa or pathways were related to chronic HF and NICM severity. The severity metrics included physician-adjudicated NYHA functional class, patient-reported KCCQ-23 scores, echocardiographic parameters, and circulating cardiac biomarker N-terminal pro-brain natriuretic peptide (NT-proBNP). Additionally, we investigated whether patients with a poor clinical outcome had unique gut microbiome signatures. Genus-level taxonomic and functional pathway data results are discussed below; Table S5 provides results on all taxonomic ranks and predicted metabolites.

After adjusting for demographics, BMI, and diet, multiple genera were associated with individual metrics of HF severity and poor clinical outcome, with a few genera associated with multiple parameters (Fig. 4a). *Butyricimonas* was associated with poor clinical outcome (*p*=0.006, *p*_adj_=0.143; Fig. 4b) and higher estimated right ventricular systolic pressure (RVSP) (*p*=0.011, *p*_adj_ = 0.116). *Longibaculum* was associated with higher LVEF (*p*<0.001, *p*_adj_=0.014; Fig. 4c), less LV enlargement (*p*=0.003, *p*_adj_=0.066), and lower RVSP (*p*=0.017, *p*_adj_=0.116). *Bifidobacterium* was associated with milder cardiomyopathy, with less LV enlargement (*p*=0.005, *p*_adj_=0.066) and less RV dysfunction (*p*=0.003, *p*_adj_=0.083). *Bifidobacterium* was also negatively associated with NYHA functional class, though only at nominal significance (*p*=0.042, *p*_adj_>0.3).

**Figure 4:**
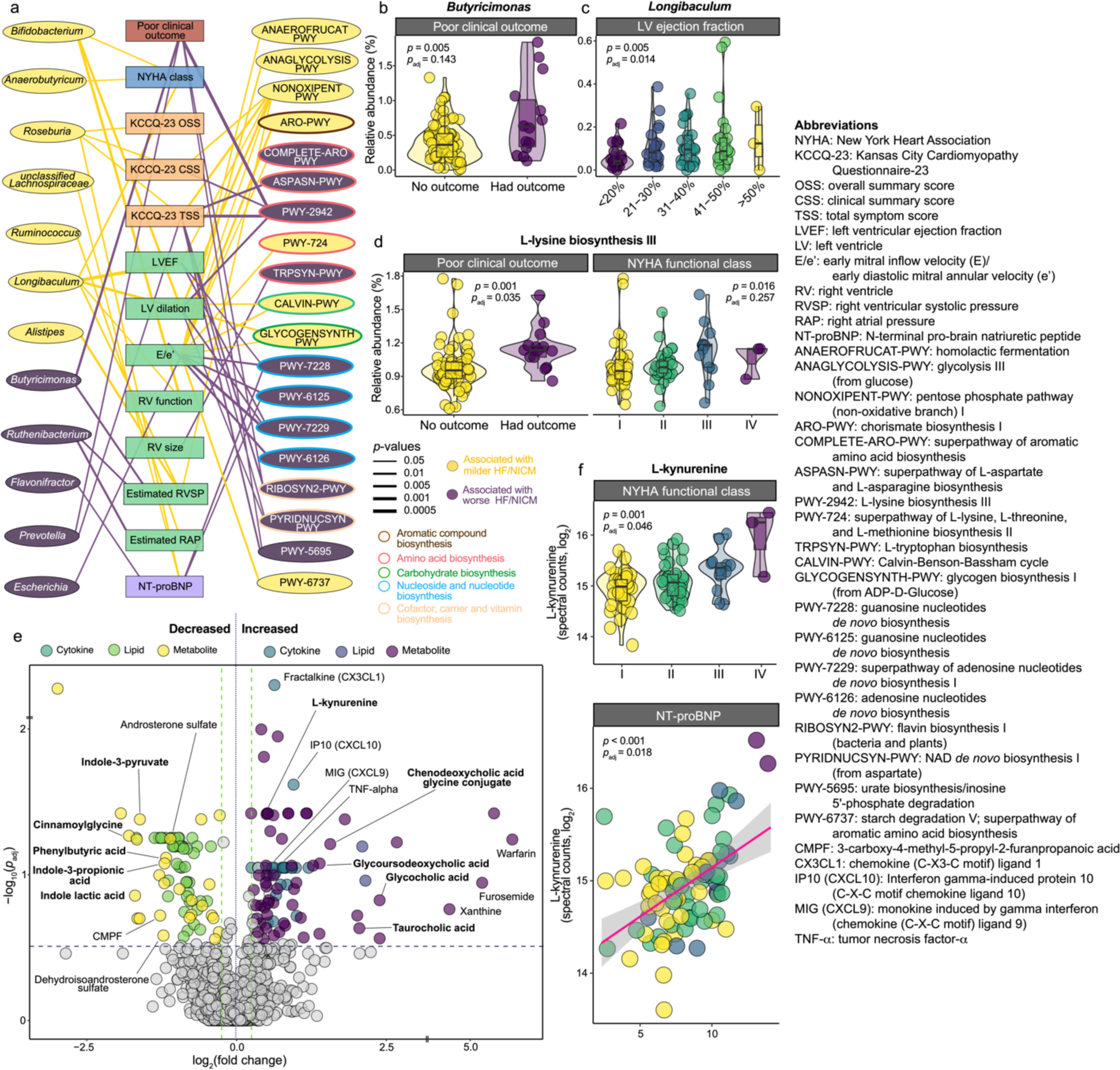
Heart failure and cardiomyopathy severity metrics are associated with gut microbiome taxa and metabolic pathways, and with circulating microbial metabolites. **a**, Several gut microbiome taxa, and metabolic pathways are associated with various metrics of disease severity, including New York Heart Association (NYHA) functional class, Kansas City Cardiomyopathy Questionnaire (KCCQ-23) scores, echocardiographic measurements, and N-terminal pro-brain natriuretic peptide (NT-proBNP), as well as risk of poor clinical outcome (a composite of death or hospice, heart transplant or left ventricular assist device (LVAD) implant). **b**, Genus *Butyricimonas* was significantly higher in individuals who had a poor clinical outcome during follow-up (*n*=12). **c**, Genus *Longibaculum* was associated with several metrics of milder heart failure and cardiomyopathy phenotype, including left ventricular ejection fraction (LVEF). **d**, L-lysine biosynthesis III (PWY-2942) was associated with several metrics of more severe HF and cardiomyopathy phenotype, including NYHA functional class, and it was independently associated with poor clinical outcome. (**a-d**), Each gut microbiome feature was modeled as a dependent variable, with each of the metrics of HF severity modeled individually as an independent variable in a linear mixed model which accounted for age, sex, race (white vs other), body mass index (BMI) and dietary patterns as fixed effects, and subject as random effect. **e**, Volcano plot of circulating cytokines, metabolites, and lipids associated with a poor clinical outcome. Many of the metabolites shown are produced by the gut microbiota, including L-kynurenine which was significantly associated with several metrics of worse HF severity, including NYHA functional class and level of NT-proBNP (log_2_), as shown in f. *p*-values were derived from the multivariate linear mixed model described for the gut microbiome features. In **e**, the log_2_ fold change was calculated for mean values of each feature by group assignment (poor clinical outcome vs no clinical outcome). (**a-f**) Benjamini-Hochberg method was used to adjust for multiple hypothesis testing.

In our microbiome functional analysis, L-lysine biosynthesis III pathway (PWY-2942) was associated with multiple metrics of HF severity (Fig. 4d), including poor clinical outcome (*p*=0.001 *p*_adj_=0.035), worse NYHA class (*p*=0.017, *p*_adj_=0.257) and KCCQ total symptom score (TSS, *p*=0.002, *p*_adj_=0.063), and more LV enlargement (*p*=0.01, *p*_adj_=0.135; Fig. S4a). Pathways for *de novo* biosynthesis of adenosine and guanosine nucleotides were also associated with worse HF severity, while the non-oxidative pentose phosphate pathway was associated with milder HF (Fig. 4a, S4b). Altogether, these findings suggest that taxonomic and functional gut microbial signatures of NICM and HF severity exist, although it remains to be determined if these are solely biomarkers of HF severity or if they contribute to chronic HF pathophysiology.

### Gut microbiome metabolites are associated with chronic HF severity

Chronic HF is a state of immune, neurohormonal, and metabolic dysregulation^30,37,38^. Thus, we next investigated how the levels of circulating cytokines, metabolites, and lipids relate to metrics of HF severity. Forty-three of 76 assayed cytokines were associated with at least one metric of HF severity (Table S6), including pro-inflammatory TNF-⍺, Interferon γ-induced Protein 10 (IP-10) and Monokine Induced by Interferon γ (MIG), consistent with previous studies^30,39–41^.

Multiple circulating metabolites were also significantly associated with chronic HF severity (Table S6; Fig. 4e, S4c,d highlight key metabolites). Notably, several of the metabolites with significant association with HF disease severity and future outcomes were metabolites known to be produced by the gut microbiota, including L-kynurenine (Fig. 4f, S4c,d), indole-3-propionic acid (IPA), phenylbutyric acid and indole lactic acid (ILA) (Fig. 4e, S4c,d). L-kynurenine was associated with several metrics of more severe HF, with other metabolites associated with milder HF. Several secondary bile acids, closely related to gut microbial metabolism, were also associated with more severe HF phenotype. Sex hormone levels (e.g., androsterone sulfate, dehydroisoandrosterone sulfate) were higher in patients with milder HF phenotype. These findings corroborate previously identified chronic HF-related bile acid dysregulation^42,43^ and catabolic state characterized by sex hormone deficiency^44–46^. Several metabolites of cardiac medications (e.g., carvedilol, losartan, furosemide, and warfarin) were also detected and associated with disease severity in predictable patterns. Circulating lipids were similarly dysregulated in individuals with chronic HF and associated with various metrics of disease severity (Table S6), although the overall magnitude of association was lower compared to circulating metabolites and cytokines.

### *Bifidobacterium* abundance is independently associated with NYHA class improvement over time

Using the longitudinal sampling of 26 subjects in our HF cohort, we investigated changes in the gut microbiome over time. Microbiome beta diversity was not significantly different between baseline and follow-up visits (*p*>0.05 for both genus and pathway comparisons) (Fig. S5a; Table S7). Instead, samples from the same subject clustered together. After accounting for multiple covariates and adjusting for multiple hypothesis testing, there were no statistically significant temporal cohort-level shifts in individual taxonomic or pathway features (Table S7). These results indicate cohort-level, between-visit gut microbiome stability in patients with chronic HF. Similar results were seen with host multi-omic data (Table S7).

While we found no significant cohort-wide gut microbiome changes, individual patients experienced divergent longitudinal microbiome trajectories. We hypothesized that these patterns might track with distinct clinical trajectories. Of the 26 patients who had multiple visits, 6 experienced improvement in NYHA class, 11 had LVEF improvement, and 8 had improved KCCQ-23 OSS. Interestingly, only 2 of these patients experienced improvement in all 3 metrics (Fig. S5b). *Bifidobacterium* was associated with improved NYHA functional class (*p*=0.001, *p*_adj_=0.029; Fig. 5a). Patients who experienced LVEF improvement had less gut *Desulfovibrio* (*p*=0.005, *p*_adj_=0.1) and *Roseburia* (*p*=0.007, *p*_adj_=0.1) (Fig. S5c). There were no significant associations with improved KCCQ-23 OSS. Table S8 lists all associations between HF improvement metrics and microbiome features.

**Figure 5:**
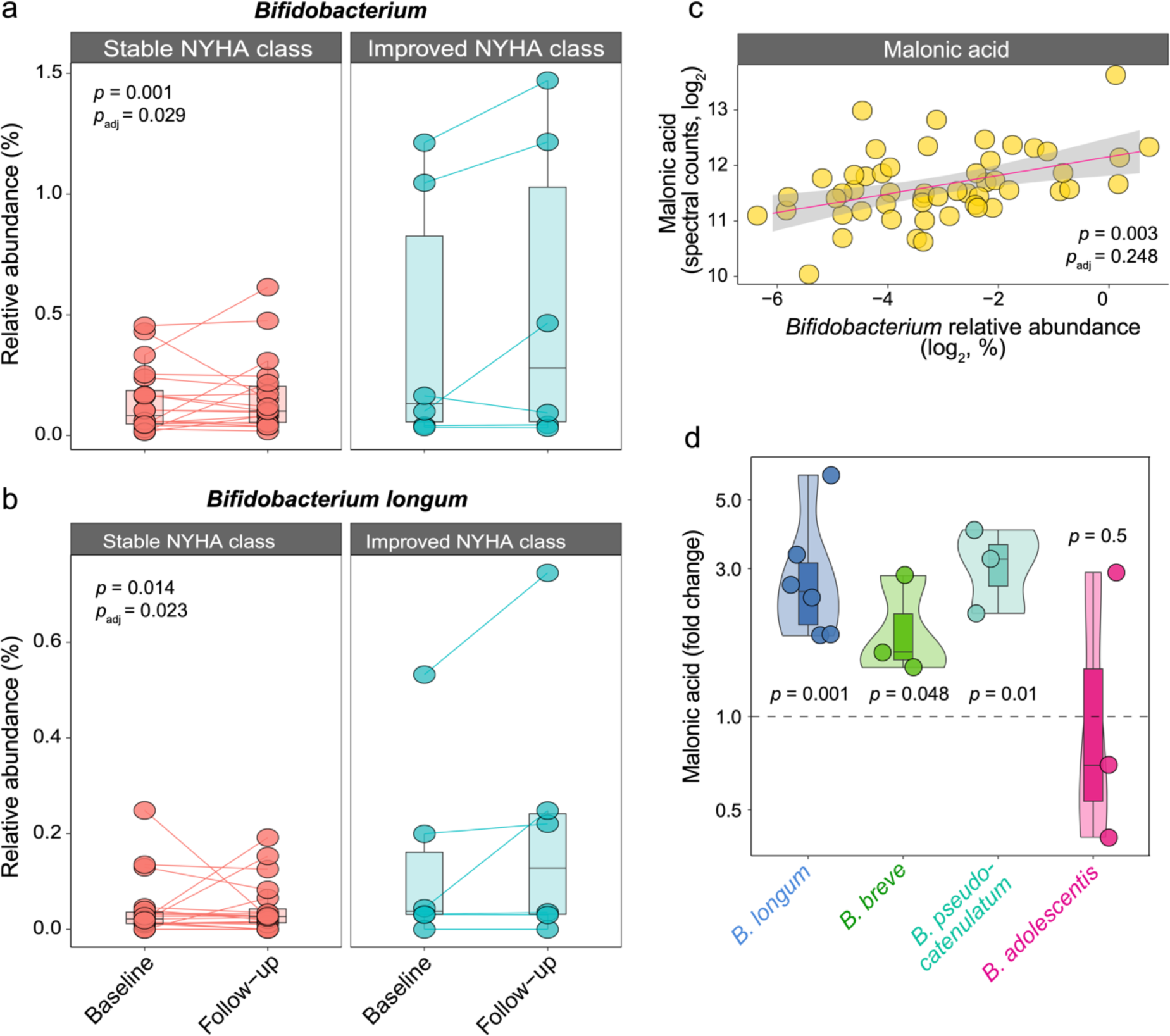
B*i*fidobacterium is associated with symptomatic HF improvement over time. **a**, Gut microbiome *Bifidobacterium* is more abundant at baseline and increases over time in individuals whose HF functional status improved over time (*n*=6, defined as an improvement in at least one class in the New York Heart Association (NYHA) functional class between baseline and follow-up visits), compared to the individuals who remain stable or worsen over time (*n*=20); **b**, Similar pattern was seen with *Bifidobacterium longum*, the most abundant of the *Bifidobacterium* species in our HF cohort; **c**, Circulating malonic acid was directly associated with the abundance of *Bifidobacterium* in the gut (shown as log_2_-transformed relative abundance for visualization). Linear mixed model with age, sex, race (white vs other), body mass index (BMI) and dietary patterns as fixed effects and subject as random effect was employed in **a-c**, followed by the Benjamini-Hochberg adjustment. In **a-b**, categorical NYHA improvement was modeled as the independent variable, with genus *Bifidobacterium* and species *B. longum*, respectively, modeled as dependent variable. In **c**, *Bifidobacterium* relative abundance was modeled as an independent variable, and circulating malonic acid as dependent. **d**, Malonic acid was produced by three out of the four *Bifidobacterium* species present in our dataset. Exometabolomics data for *B. longum*, *B. breve*, *B. pseudocatenulatum*, and *B. adolescentis* were extracted from the published dataset^47^. Fold changes were calculated in reference to the base media (Mega Media), with *p*-values computed from a one-tailed Student’s t-test with *n*=6 for *B. longum*, and *n*=3 for other species.

Fifty-one patients had long-term clinical follow-up data available (average follow-up of 27 months). Long-term improvement in NYHA (*n*=7) was directly associated with *Bifidobacterium* genus (*p*=0.005, *p*_adj_=0.138; Fig. S5d) and *Veillonellaceae* family (*p*=0.013, *p*_adj_=0.11), while negatively associated with *Rikenellaceae* family (*p*=0.034, *p*_adj_=0.191). Long-term LVEF improvement (*n*=10) was not associated with any microbiome signatures (Table S8).

Having identified genus *Bifidobacterium* as significantly associated with functional HF improvement, we investigated how the four *Bifidobacterium* species detected in our cohort related to longitudinal changes in HF clinical status. Due to their low relative abundance in the HF cohort, these species were not initially included in our species-level taxonomic analysis. Three of these species (*Bifidobacterium longum* (Fig. 5b), *B. breve*, and *B. pseudocatenulatum*) were significantly associated with NYHA improvement from baseline to follow-up visit (*p*=0.013, *p*=0.014, *p*=0.018, respectively; *p*_adj_=0.024 for all; Table S9). These species were also associated with several HF severity metrics and indicators of milder HF and NICM (Table S9). Altogether, these analyses suggest *Bifidobacterium* as a biomarker of clinical HF improvement.

### *Bifidobacterium* is associated with multiple circulating host molecules

We next sought to understand the relationship between *Bifidobacterium* abundance and the host circulating metabolome (*n*=566 features), lipidome (*n*=737), and immunome (*n*=76), to probe potential mechanisms through which *Bifidobacterium* may affect HF progression. Multivariate analyses showed that *Bifidobacterium* abundance was directly associated with circulating malonic acid (*p*=0.002, *p*_adj_=0.248; Fig. 5c), and inversely associated with circulating lysophosphatidylethanolamine (LysoPE-22:6, *p*=0.001, *p*_adj_=0.248) and lysophosphatidylcholine (LysoPC-22:6, *p*=0.003, *p*_adj_=0.248) (Fig. S5e). Although only at nominal significance, *Bifidobacterium* abundance was also positively associated with circulating IPA (*p*=0.021) and ILA (*p*=0.043) (Fig. S5e), and many lipids (Table S10).

### *Bifidobacterium* species produce malonic acid *in vitro*

To further explore the positive association between *Bifidobacterium* and circulating malonic acid within our chronic HF cohort, we investigated whether *Bifidobacterium* species can produce this metabolite. A previous exometabolomics study^47^ found that *B. longum*, *B. breve*, and *B. pseudocatenulatum* produced and secreted malonic acid (Fig. 5d). The first three species also produced ILA, although none produced IPA. These data suggest that *Bifidobacterium* species synthesize and potentially contribute malonic acid into the host circulation, with conceivable downstream effects on host physiology that may be relevant in the context of chronic HF.

## DISCUSSION

In this longitudinal, multi-omic study of patients with NICM-related systolic HF, we identified significant differences in gut microbiome taxonomic and functional composition and differential connections between the gut microbiome and the host immune system in patients with chronic HF compared with healthy subjects. Furthermore, we uncovered gut microbiome signatures of chronic HF severity and identified *Bifidobacterium* as a biomarker of HF clinical improvement (Fig. 6).

**Figure 6:**
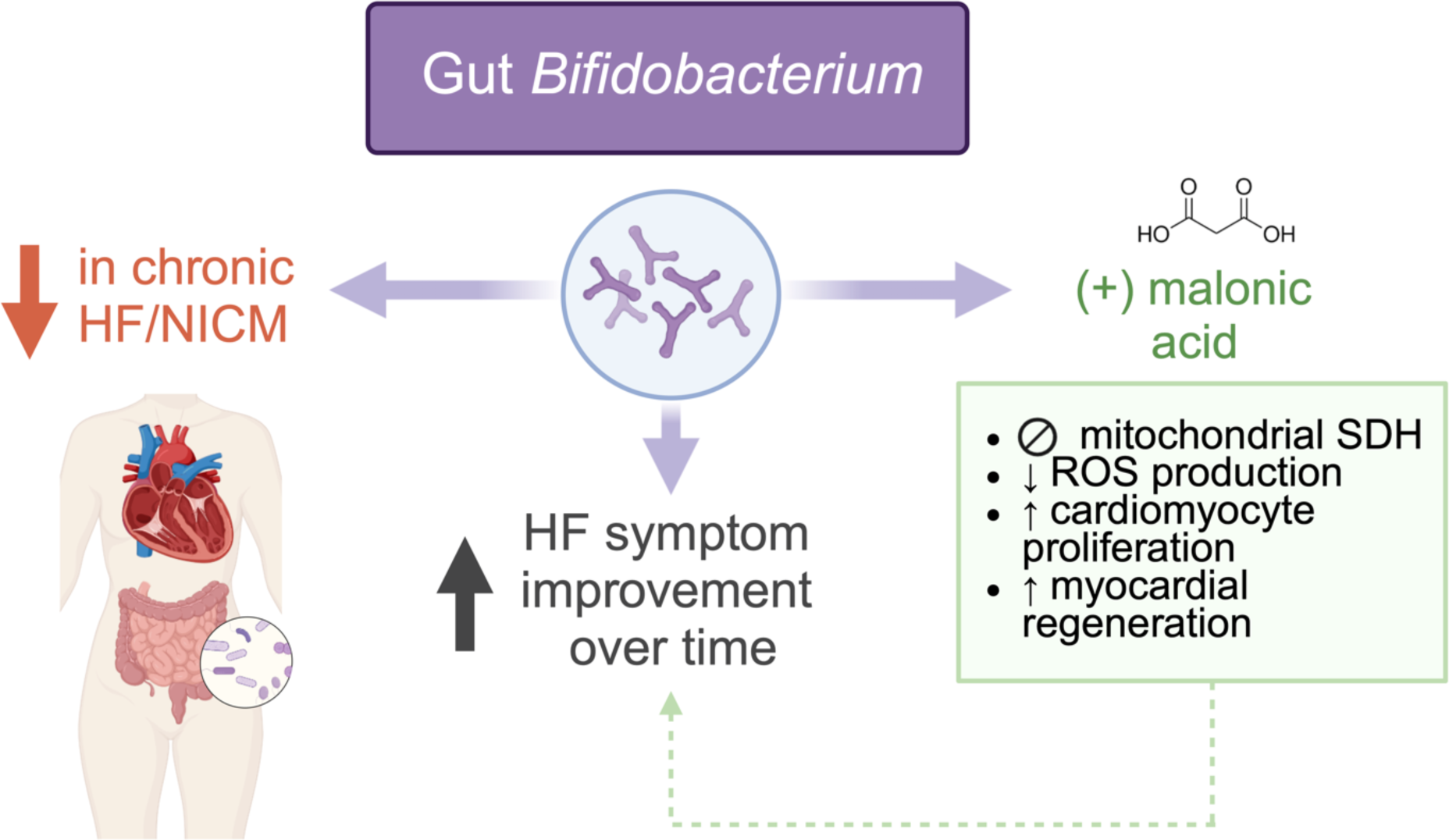
Summary of study findings. Gut microbiome *Bifidobacterium* is depleted in adults with non-ischemic cardiomyopathy (NICM)-related chronic heart failure (HF). This microbe is also associated with HF improvement over time (expressed as between-visit change in the New York Heart Association functional class). *Bifidobacterium* is also directly associated with circulating malonic acid, which is produced by several *Bifidobacterium* species. Malonic acid has been implicated in biological processes dysregulated in chronic HF. Abbreviations: SDH: succinate dehydrogenase; ROS: reactive oxygen species. (Figure created with BioRender.com.)

Our findings that the chronic HF-associated gut microbiome is significantly altered compared to healthy subjects, with decreased alpha diversity and a profound loss of anti-inflammatory gut microbes, further validates earlier studies of gut microbiome in chronic HF^11–13^. Interestingly, data on *Bifidobacterium* in chronic HF is mixed: two small studies found *Bifidobacterium* enrichment in patients with decompensated^48^ and severe HF^49^, but several larger studies support our finding of *Bifidobacterium* depletion^6,11^. We also identified enrichment in certain pro-inflammatory gut microbes and potential pathogens, which has been postulated to contribute to the gut barrier compromise in chronic HF, thus driving low-grade inflammation and disease progression^5,21,22^.

Several of the microbes that were depleted in the HF cohort are SCFA producers, consistent with the depletion of SCFA fermentative pathways. The cardiovascular effects of SCFAs are diverse and profound, including regulation of blood pressure, cardiac hypertrophy, cardiorenal fibrosis, and post-injury repair^14–18^. Enrichment of pathways for fatty acid beta oxidation suggest the presence of an abnormally oxygen-rich environment within the gut and a shift away from anaerobic fermentation. Such a shift facilitates the expansion of facultative anaerobes, many of which are pro-inflammatory and/or pathogenic^18,27,28^. Moreover, the predicted increase in formaldehyde burden within the gut lumen due to depletion of formaldehyde detoxification pathways and related enrichment of folate biosynthetic pathways (major sources of formaldehyde) may lead to oxidative toxicity and inflammation of the host gut epithelium^50^.

Our multi-omic machine learning model, which combined clinical laboratory data with host cytokine and gut microbiome data, was excellent at classifying chronic HF compared to healthy state. Moreover, the model based only on microbiome pathways had nearly identical classification ability, as did the model that included only top 50 distinguishing features. Of these, gut microbiome features represented a substantial majority, comprising pathways related to SCFA biosynthesis and formaldehyde detoxification, as well as genera *Bifidobacterium*, *Longibaculum*, and other SCFA producers. In general, our findings suggest that microbiome features (especially functional ones) could be used as diagnostic biomarkers for HF, although they should be validated in larger, independent chronic HF cohorts with a wider range of disease severity, and likely against other comorbidities given the shared microbiome dysbiosis among conditions associated with low grade inflammation (e.g., diabetes)^10,51^.

Our study also identified several gut microbiome features associated with chronic HF severity. *Butyricimonas* was associated with higher risk of poor clinical outcome. Although *Butyricimonas* members are butyrate producers and were found to be related to insulin sensitivity and lower BMI^52,53^, other studies identified *Butyricimonas* enrichment in patients with coronary artery disease^54^ and irritable bowel syndrome^55^. *Longibaculum*, which was significantly depleted in our chronic HF cohort and associated with a milder HF phenotype, was previously associated with improved metabolic health^56–58^ and responsiveness to supplementation with fiber^59^ and probiotics^60^. Thus, it is possible that some of the positive metabolic effects of *Longibaculum* can beneficially affect HF pathophysiology. Gut microbiome L-lysine biosynthesis was enriched in patients with HF compared to healthy individuals and associated with several indicators of worse HF severity and future risk of poor clinical outcome. These findings align with previous studies that found L-lysine metabolism was dysregulated in a rat model of pressure-overload hypertrophy and HF^61^.

Circulating gut microbiome-produced metabolites showed strong association with chronic HF severity. Downstream microbial metabolites of dietary tryptophan IPA and ILA were associated with milder HF. IPA modulates mitochondrial function in cardiomyocytes^62^, protects from diastolic dysfunction in the context of HF with preserved LVEF^63^, and ameliorates doxorubicin cardiotoxicity^64^. Less is known about the cardiovascular effects of ILA, although it is known to strengthen gut barrier function and plays an anti-inflammatory role^65–67^. On the contrary, L-kynurenine, another gut microbiome-produced tryptophan metabolite, was associated with more severe HF, consistent with previous studies linking kynurenine-related metabolites to HF severity and worse outcomes^68–70^.

Integration of our longitudinal clinical and microbiome data revealed *Bifidobacterium* enrichment in patients whose functional status improved over time, a finding corroborated by another study^71^. While *Bifidobacterium* abundance may be solely a biomarker of clinical HF improvement – a consequence of gut milieu changes with HF improvement – significant associations with host circulating metabolites that have been implicated in cardiac and HF physiology suggest that *Bifidobacterium* species may directly (or indirectly) contribute to HF improvement. For example, *Bifidobacterium* was closely associated with circulating malonic acid. Malonic acid is a reversible inhibitor of succinate dehydrogenase (SDH), a key enzyme that links Krebs cycle and mitochondrial respiratory chain. After myocardial injury, SDH oxidizes the excess mitochondrial succinate with resultant reactive oxygen species production and mitochondrial injury and cell death^72^ – all of these downstream effects are ameliorated by malonate administration^73^. In another animal model, malonate-mediated SDH inhibition, after an ischemic insult, promoted adult cardiomyocyte proliferation and myocardial regeneration through a metabolic shift from oxidative phosphorylation to glucose metabolism^74^. Beyond these direct myocardial effects, malonate was also shown to suppress the p38 MAPK/NFkB pathway and thus LPS-induced tissue inflammation^75^. Our data suggest that several *Bifidobacterium* species can produce malonic acid, thus potentially contributing to the host circulating malonic acid pool, which could exert systemic and localized target organ effects, including the heart, through these pathways. Future clinical studies should evaluate if *Bifidobacterium* probiotic supplementation would directly alter HF physiology and lead to clinical improvement. These studies could also determine if the mechanisms that we have proposed (e.g., production of metabolites such as malonic acid) mediate the effects of *Bifidobacterium* species on the host.

### Strengths

This is the first study of the chronic HF-associated gut microbiome that combines longitudinal design, metagenomic gut microbiome sequencing (which allows for both taxonomic and functional microbiome assessment), and multi-faceted and detailed host clinical and sub-clinical multi-omic data. This combination of data enabled us to deeply dissect chronic HF-associated gut microbiome composition and its links with the host immune system. Additionally, the longitudinal data, combined with clinical assessments, enabled us to identify microbiome signatures associated with HF clinical trajectories. Furthermore, by integrating host multi-omic data with gut microbiome profiles, we identified host molecules through which the gut microbiome may affect the host and modulate HF physiology. We also collected dietary data, which allowed us to correct for the effect of dietary habits, as well as detailed data on patients’ medications and comorbidities.

### Limitations

Our study has multiple limitations. This was a single-center study in which there was demographic imbalance (e.g., both cohorts were predominantly white, and men were overrepresented in the HF cohort). Due to a between-cohort technical batch effect, untargeted metabolomic and targeted lipidomic data could not be compared between the HF and healthy cohorts. There were also differences in the method of stool sample collection between the HF and healthy cohorts. However, our analysis of paired stool samples from healthy individuals using both methods showed no significant batch effects (Methods). The results of our study are not necessarily applicable to all patients with HF, as our study included only subjects with NICM-related HF with reduced, mildly reduced, and recovered LVEF, and excluded subjects with treated diabetes and other serious comorbidities. Furthermore, our cohort was relatively small and clinically well-compensated as a whole, which likely skewed our results. Even so, we were able to detect many gut microbiome and multi-omic signals associated with HF severity and clinical change.

### Summary

This is the first longitudinal gut microbiome multi-omic study of patients with NICM-related chronic HF, in which we describe HF-associated taxonomic and functional gut microbiome signatures. We also identify gut microbiome signatures associated with HF disease severity. Furthermore, we find that *Bifidobacterium*, severely depleted in patients with chronic HF, is directly related to clinical HF improvement over time, and associated with host circulating malonic acid. We also show that multiple *Bifidobacterium* species produce malonic acid *in vitro*. We thus pinpoint a plausible mechanism through which *Bifidobacterium* may impact HF physiology and modulate disease progression. Future clinical studies of *Bifidobacterium* probiotic supplementation should be pursued to evaluate if *Bifidobacterium* is merely a biomarker of clinical HF improvement or an active contributor to cardiac physiology.

## METHODS

### Study population

#### Patients with non-ischemic cardiomyopathy (NICM)/chronic heart failure (HF)

Between July 2018 and March 2020, 59 adults followed at the Stanford Cardiomyopathy Clinic were enrolled in the study. Participants provided written informed consent for the study under research protocol 44967 approved by the Stanford University Institutional Review Board. We included adults (ages 18-75) with established diagnosis of NICM and reduced left ventricular ejection fraction (LVEF) (LVEF < 50%) as determined by transthoracic echocardiogram. We excluded individuals with primary ischemic cardiomyopathy or complex congenital heart disease. Other exclusion criteria included previously known and treated diabetes mellitus, advanced chronic kidney disease (stage 4 or 5), advanced liver disease or cirrhosis, systemic autoimmune or connective tissue disorder, active malignancy, substantial gastrointestinal disease (including inflammatory bowel disease, celiac disease, severe chronic diarrhea or constipation), any prior history of bowel resection, history of appendectomy or cholecystectomy in the preceding 6 months, or chronic conditions requiring systemic immunosuppressive or immunomodulatory agents. We also excluded individuals who reported receiving systemic antibiotics or probiotics in the preceding month, or systemic chemotherapy or radiation therapy in the preceding 12 months.

#### Healthy subjects

Subjects who served as healthy controls for our study were healthy adults who had previously been recruited for the integrated Personal Omics Profiling (iPOP) study, one of the core studies of the National Institutes of Health (NIH) Integrative Human Microbiome Project (iHMP)^32,76,77^. The goal of this multi-omic, longitudinal, prospective cohort study of 109 subjects was to investigate systems-level biological changes that characterize transition from a healthy to pre-diabetic or diabetic state. The study subjects were profiled extensively on a quarterly basis while healthy, with more frequent visits during acute viral illnesses, following immunizations, or for other interventions (e.g., intentional weight gain and weight loss). For inclusion in our study, subjects needed to have both stool and fasting blood samples from at least one healthy visit available. We excluded subjects diagnosed with diabetes (treated or untreated), any cardiovascular comorbidities, or other non-cardiac comorbidities outlined in the chronic HF cohort. Other exclusion criteria included treatment with systemic antibiotics or probiotics in the preceding month. Ultimately, 50 of the iPOP study subjects were included in our study. This cohort was recruited between 2010 and 2018.

### Study procedures

#### Stool samples

Stool samples were obtained from a total of 109 study subjects. Patients with chronic HF collected freshly produced stool samples using an OMNIgene-GUT collection kit (Cat. #: OMR-200). Samples were kept at room temperature for up to 15 days before transportation and were then aliquoted and frozen at −80 °C; the OMNIgene kit contains a stabilizing buffer that maintains gut microbiome profiles for up to 60 days (https://www.dnagenotek.com/row/pdf/PD-WP-00056.pdf). Healthy participants had previously collected stool samples according to the Human Microbiome Project Core Microbiome Sampling Protocol A (https://www.hmpdacc.org). Briefly, these stool samples were collected through a conventional method without a stabilization buffer with samples immediately frozen at −20 °C, transported on ice to their study visit, upon which they were placed at −80 °C for long-term storage.

#### Blood samples

Blood sample collection for patients with chronic HF followed protocols established by the iPOP study^32^. At each study visit, following an overnight fast (10-12 h), patients had a blood draw for a full panel of clinical laboratory tests, which included complete blood count, complete metabolic panel, insulin, hemoglobin A1c (HbA1c), lipid panel with direct low-density lipoprotein (LDL), and cardiac biomarker N-terminal pro-brain natriuretic peptide (NT-proBNP). Fasting blood was additionally drawn for research blood tests, including a cytokine panel, untargeted metabolomics, and targeted lipidomics. All blood draws and clinical laboratory tests were performed in the Stanford Clinical Laboratory according to standard protocols. After collection, blood samples were processed according to a previously published protocol^32^ and stored at −80 °C for future analysis.

#### Kansas City Cardiomyopathy Questionnaire-23 (KCCQ-23)

At each study visit, individuals with chronic HF were asked to complete the KCCQ-23 survey. This survey is a patient-reported outcome instrument developed to independently measure patient’s perception of their health status and has been well validated in patients with chronic HF^78^. The questionnaire is comprised of 23 questions, which are aggregated into a score. Each score ranges from 0 to 100, where lower score denotes increasing symptoms or limitations to daily functioning due to HF symptoms. Healthy subjects did not complete this survey.

#### Dietary questionnaire

Dietary questionnaire data was available for both cohorts. The dietary survey was a food frequency questionnaire (FFQ) inquiring about frequency of intake of most common food items and was modified from the Scottish Collaborative Group FFQ (https://www.foodfrequency.org/). The questionnaire asked about habitual consumption of 22 dietary items. Out of 109 subjects and 136 visits, data from 2 visits by individuals with chronic HF and 4 visits by healthy individuals were not available.

Some dietary data were missing from 26 visits (mean 10.1%, standard deviation 10.0). Missing data were imputed using the *k*-nearest neighbors method function (“impute.knn”) in the R “impute” package (v. 1.46.0). Due to a large number of dietary variables relative to the number of samples, we aggregated individual dietary items into 6 food categories (e.g., “fiber” food category was calculated as a sum of reported intake of beans and lentils, green vegetables, root vegetables, salad, and fresh fruit).

#### Clinical data collection

Demographic data and clinical data on cardiac status, relevant cardiac history and echocardiographic imaging, poor clinical outcomes (being listed for or receiving a heart transplant, implantation of a durable left ventricular assist device (LVAD), placement on hospice for end-stage HF, or death), relevant comorbidities, and prescribed medications were extracted from the electronic medical record (EMR) from the HF clinic visits coinciding with the study visit.

## Multi-omic measures and data processing

### Metagenomic sequencing

#### Library preparation and sequencing

Genomic DNA was extracted from stool using a QIAamp PowerFecal DNA Kit (Cat. #: 12830-50). Libraries were prepared according to a standard Illumina protocol, using a TruSeq DNA PCR-Free Low Throughput Library Prep Kit (SKU 20015962) with 1 mg of input DNA. Paired-end sequencing was performed on a HiSeqX at Psomagen, Inc. (Rockville, MD). A total of 3,930,771,952 paired-end 150-bp reads were acquired for all samples, with an average of 17,548,089 reads per sample. Samples were sequenced in 2 batches, with the first batch using all 8 lanes of a flow cell. Samples with a low number of reads were separately re-sequenced in a single flow-cell lane. Multiple control samples were also sequenced in various lanes in the first batch, as well as in the second batch. There was no clear bias between the two sequencing batches. For each sample, only the sequencing run with the most reads was used for downstream analysis.

#### Metagenome quality control and deduplication

Raw sequencing reads were demultiplexed. Adapters and low-quality bases were trimmed using Trimmomatic^79^ with default parameters. Reads from human cells were removed using KneadData (https://github.com/biobakery/kneaddata) against the human genome (hg37 reference database). Exact duplicate reads were removed with clumpify (BBtools) (https://sourceforge.net/projects/bbmap/).

#### Estimating species and pathway abundances from metagenomic data

Bracken v. 2.6^80^ was used to estimate taxonomic abundances from metagenomic reads. Each read was first assigned a taxonomic classification using exact k-mer matches with Kraken v. 2.0^81^. Final taxonomic abundance was re-estimated using a Bayesian method with Bracken. The abundances of microbial metabolic pathways and gene families were profiled using HUMAnN v. 3.0^82^ with default parameters. Microbiome features that were estimated to have relative abundance of <0.01% within a sample were assumed to be false positive reads and were re-assigned a relative abundance of 0%. The relative abundance cut-off for microbial pathways, due to their overall lower individual relative abundance, was <0.001%. Taxa and pathways were additionally filtered out if they were not present in at least half of all analyzed samples. The remaining taxa and pathways were included in HF versus healthy analyses.

For analyses focused on the HF cohort, taxa and pathways were further filtered. We hypothesized that the gut microbial features with the strongest influence on host biology in chronic HF would be those that are either differentially abundant between the HF and healthy cohorts, or present at high relative abundance within the HF cohort (top 10 most abundant taxa that are not represented in the differentially abundant group for family, genus, and pathway levels, and top 25 most abundant microbial species given their lower relative abundance on this taxonomic level). Thus, our HF analyses included only these limited features (17 families, 26 genera, 36 species, and 31 pathways).

#### Functional enrichment analysis

The metabolic pathways identified by HUMAnN v. 3.0 were hierarchically classified based on the MetaCyc database^83^. The Gene Set Enrichment Analysis (GSEA) method^26^ was applied to each classification level to identify pathway groups with significantly different levels between the cohorts.

#### Predicting metabolite composition from metagenomic data

Microbial metabolite compositions in stool samples were predicted using MelonnPan^29^ with its pre-trained model from the human gut microbiome using the gene family estimates from HUMAnN v. 3.0. Of the 107 metabolites that MelonnPan predicts in its original validation dataset, 80 (75%) were identified in our metagenomics data.

### Immune protein measurements

#### Sample preparation

Fasting serum samples were prepared as per the iPOP study protocol^32^ and stored at −80 °C for future analysis.

#### Data acquisition

Levels of circulating cytokines in blood were measured using a 76-plex Luminex antibody-conjugated bead capture assay (EMD Millipore) that has been extensively characterized and benchmarked by the Stanford Human Immune Monitoring Center (HIMC)^84^. The assay was performed by the HIMC-Immunoassay Team. Kits were used according to the manufacturer’s recommendations with modifications as follows. The kits include three panels. Panel 1 is Milliplex HCYTMAG-60K-PX41 with the addition of IL-18 and IL-22. Panel 2 is Milliplex HCP2MAG-62K-PX23 with the addition of MIG. Panel 3 includes Milliplex HSP1MAG-63K-06 and HADCYMAG-61K-03 (RESISTIN, leptin, and HGF) to generate a 9-plex. Samples were diluted 3-fold for panels 1 and 2 and 10-fold for panel 3. Diluted sample (25 µL) was mixed with antibody-linked magnetic beads in a 96-well plate and incubated overnight at 4 °C with shaking.

Cold and room temperature incubation steps were performed on an orbital shaker set at 500-600 rpm. Plates were washed twice with a wash buffer in an ELx405 washer (BioTek Instruments). Following a 1-h incubation at room temperature with biotinylated detection antibody, streptavidin-PE was added for 30 min with shaking. Plates were washed as described above and PBS was added to wells for reading in a Luminex FlexMap3D with a lower bound of 50 beads per sample per cytokine. Custom Assay Chex control beads (CUSTOM ACX) were added to all wells (Radix BioSolutions, Georgetown, Texas). Wells with a bead count <20 were excluded.

#### Data processing

Samples from the two cohorts (chronic HF and healthy) were randomly distributed between the two plates but repeat samples from the same subject were analyzed on the same plate. Inter-plate variability was corrected using the median of inter-plate ratios for four representative samples in each plate. Raw mean fluorescence intensity values were used for the analysis. Missing values (bead count <20) were imputed using the *k*-nearest neighbors method function (“impute.knn”) in the R “impute” package (v. 1.46.0). Cytokines were log_2_-transformed prior to downstream analyses.

### Untargeted metabolomics and targeted lipidomics

#### Sample preparation

Plasma samples were thawed on ice, prepared, and analyzed randomly as previously described^85^. Metabolites and complex lipids were prepared from 40 µL of plasma using a biphasic separation with cold methyl tert-butyl ether, methanol, and water and reconstituted in 9:1 methanol:toluene. Each sample was spiked-in with 15 labeled metabolite internal standards (IS) to control for extraction efficiency and liquid chromatography-mass spectrometry (LC-MS) performance and deuterated lipid IS (Sciex, Cat. #: 5040156, lot #: LPISTDKIT-102b) for quantification. For quality controls, 3 reference plasma samples (40 µL) and 1 preparation blank were processed in parallel.

#### Data acquisition

Plasma metabolites were analyzed using a broad-spectrum metabolomics platform consisting of hydrophilic interaction chromatography (HILIC) and reverse phase liquid chromatography (RPLC)–MS^86^. Complex lipids were quantified using a targeted MS-based approach^85^.

<u>1) Untargeted metabolomics by LC-MS.</u> Metabolic extracts were analyzed 4 times using HILIC and RPLC separation in both positive and negative ionization modes. Data were acquired on a Thermo Q Exactive HF mass spectrometer for HILIC and a Thermo Q Exactive mass spectrometer for RPLC operated in full MS scan mode. MS/MS data were acquired on quality control samples (QC) consisting of an equimolar mixture of all samples in the study. HILIC experiments were performed using a ZIC-HILIC column (2.1 × 100 mm, 3.5 μm, 200 Å; Merck Millipore, Darmstadt, Germany) and mobile phase solvents consisting of 10 mM ammonium acetate in 50/50 acetonitrile/water (A) and 10 mM ammonium acetate in 95/5 acetonitrile/water (B). RPLC experiments were performed using a Zorbax SBaq column (2.1 × 50 mm, 1.7 μm, 100 Å; Agilent Technologies, Palo Alto, CA) and mobile phase solvents consisting of 0.06% acetic acid in water (A) and 0.06% acetic acid in methanol (B). Data quality was ensured by (i) injecting 6 and 12 pool samples to equilibrate the LC-MS system prior to running the sequence for RPLC and HILIC, respectively; (ii) injecting a pool sample every 10 injections to control for signal deviation with time; and (iii) checking mass accuracy, retention time, and peak shape of IS in each sample.
<u>2) Targeted lipidomics using the Lipidyzer Platform.</u> Lipid extracts were analyzed using a commercial flow injection approach namely the Lipidyzer platform^85^. Briefly, lipid molecular species were identified and quantified using multiple reaction monitoring (MRM) and positive/negative ionization switching. Two acquisition methods were employed covering 13 lipid classes; method 1 had SelexION voltages turned on while method 2 had SelexION voltages turned off. Data quality was ensured by i) tuning the DMS compensation voltages using a set of lipid standards (Cat. #: 945 5040141, Sciex) after each cleaning, more than 24 h of idling, or 3 days of consecutive use; ii) performing a quick system suitability test (QSST) (Cat. #: 50407, Sciex) before each batch to ensure an acceptable limit of detection for each lipid class; and iii) triplicate injection of lipids extracted from a reference plasma sample (Cat. #: 948 4386703, Sciex) at the beginning of each batch.

#### Data processing

<u>1) Untargeted metabolomics.</u> Data from each mode were independently processed using Progenesis QI software v. 2.3 (Nonlinear Dynamics, Durham, NC). Metabolic features from blanks and those that did not show sufficient linearity upon dilution in QC samples (*r*<0.6) were discarded. Only metabolic features present in >2/3 of the samples were kept for further analysis. Inter- and intra-batch variations were corrected using systematic error removal with random forest normalization method on QC injected repetitively along the batches^87^. Missing values were imputed by drawing from a random distribution of low values in the corresponding sample. Data from each mode were merged and resulted in a dataset containing 5681 metabolic features that was used for downstream analysis. Metabolic features of interest were tentatively identified by matching fragmentation spectra and retention time to analytical-grade standards (MSI=1) when possible or matching experimental MS/MS to fragmentation spectra (MSI=2) in publicly available databases, resulting in total of 566 individual metabolites. Metabolite abundances were reported as spectral counts. Raw and processed data will be made available on the Metabolomics Workbench data repository before manuscript is published. Prior to analyses, each metabolite abundance was log_2_-transformed.
<u>2) Targeted lipidomics.</u> Lipidyzer data were reported by the Lipidomics Workflow Manager software, which calculates concentrations for each detected lipid as the average intensity of the analyte MRM/average intensity of the most structurally similar IS MRM multiplied by its concentration. Lipids detected in <2/3 of the samples were discarded, and missing values were imputed by drawing from a random distribution of low values class-wise in the corresponding sample. A dataset containing 737 individual lipid species was used for downstream analysis. Lipid abundances were reported as concentrations in nmol/g, which were then log_2_-transformed prior to analyses.

### In vitro data

#### *In vitro* exometabolomics data for *Bifidobacterium* species

Exometabolomics data for *B. longum*, *B. breve*, *B. pseudocatenulatum*, and *B. adolescentis* were extracted from a previously published dataset^47^. We only included measurements in which cells grew to early stationary phase and excluded data from cultures with poor growth (OD < 0.1). Fold changes were calculated in reference to the base medium (Mega Media). *p*-values were computed from a one-tailed Student’s t-test with *n*=6 for *B. longum*, and *n*=3 for the other species.

### Statistical analyses

#### Clinical metadata

Clinical data included demographic, anthropomorphic, laboratory, and relevant cardiovascular clinical data. Continuous variables were evaluated for normal distribution using the Shapiro-Wilk test. For baseline comparison of the individuals from the HF and healthy cohorts, Student’s t-tests were used for normally distributed variables. Mann-Whitney U tests were used for continuous variables with skewed distribution, and Fisher’s exact tests were used for categorical variables.

### Metagenomic data

#### OMNIgene-GUT vs regular stool sample collection methods

Paired samples (*n*=36) were collected to evaluate sample collection method-related batch effects. A linear mixed model (LMM) was used to evaluate if microbiome features were related to the collection method, with the latter modeled as an independent categorical variable and microbial features modeled as dependent. Microbial features analyzed included alpha diversity metrics (Chao1, Shannon index, number of observed species, and Simpson diversity index), taxonomic (phylum, family, genus, and species level), and functional features (microbial pathways). Beta diversity analysis was performed on different taxonomic ranks and on microbial pathways, using UniFrac (unweighted and weighted) and Bray-Curtis distance, respectively. Permutational multivariate analysis of variance (PERMANOVA) was used for statistical testing.

#### Comparison between HF and healthy cohorts

For individual taxonomic and functional microbiome features, LMM was used that accounted for subject age, sex, race, and body mass index (BMI), as well as reported habitual dietary patterns as fixed effects, and subject as a random effect. The microbiome features (including alpha diversity metrics, taxa, microbial pathways, and MelonnPan-predicted stool metabolites^29^) were modeled individually as dependent variables, with categorical HF modeled as an independent variable. Beta diversity analysis was performed using weighted UniFrac and Bray-Curtis distance for taxa and pathways, respectively, with PERMANOVA statistical testing. For functional enrichment analysis, GSEA method^26^ was applied (described above) with *p*-values of each pathway group estimated via 1,000 permutations and adjusted for multiple hypothesis testing.

#### Within HF cohort analysis (association with chronic HF phenotype, disease severity, and poor clinical outcome)

For these analyses, restricted to the HF cohort, LMM was used that accounted for subject age, sex, race, BMI, and diet as fixed effects, with subject as a random effect. Race was modeled as ‘white’ vs ‘other’ given the overall smaller dataset (*n*=59 individuals, with 86 total visits) and relatively few individuals belonging to each individual non-white category. Microbial features were modeled as dependent variables, with HF phenotype, severity, or clinical change variable modeled as independent. The clinical metrics included physician-adjudicated New York Heart Association (NYHA) functional class, patient-reported KCCQ-23 scores (overall summary score (OSS), clinical summary score (CSS), and total symptom score (TSS)), echocardiographic parameters (LVEF, left ventricular (LV) dilation, E/e’, right ventricular (RV) dilation, RV function, estimated RV systolic pressure (RVSP), and estimated right atrial pressure (RAP)), circulating cardiac biomarker NT-proBNP (log_2_-transformed due to its distribution), and risk of future poor clinical outcome.

#### Longitudinal data for chronic HF cohort

With the availability of longitudinal data for a subset of the HF cohort (*n*=26, with 52 total visits; for one individual who recorded 3 visits, we included only their baseline and last visit), we investigated both cohort-level between-visit gut microbiome shifts, and gut microbiome signatures associated with significant between-visit HF clinical change. For the former, beta diversity analysis was performed using Bray-Curtis distance with PERMANOVA testing. For individual microbiome features, an LMM was used with multivariate adjustment described above, and visit number (baseline versus follow-up) was modeled as an independent variable.

A similar analysis was performed for significant clinical change, which was defined as: (1) between-visit improvement in documented NYHA functional class, defined as a decrease in at least 1 level (*n*=6 patients), (2) LVEF improvement, defined as ≥5% increase in LVEF (*n*=11 patients), and (3) KCCQ OSS improvement, defined as ≥5-point increase in OSS between visits (*n*=8 patients). The binary variable (‘stable’ versus ‘improved’) was modeled as an independent variable, with each microbiome feature modeled as dependent.

The majority of the cohort had long-term clinical EMR data available (*n*=51 individuals with 75 total visits, as some individuals had only one study visit) with an average follow-up interval of 27 months. The clinical data of interest included reported NYHA functional class and LVEF, with the improvement defined as: (1) decrease in NYHA class of at least 1 level, and (2) increase in LVEF of ≥5% from baseline to long-term. The statistical modeling was similar to the above longitudinal analysis.

### Multi-omic host data (cytokine, untargeted metabolomic and targeted lipidomic data)

#### Chronic HF vs healthy

Due to a cohort-related batch effect in plasma samples, only serum samples and thus cytokine data (*n*=76) could be compared between the chronic HF (*n*=58) and healthy cohort (*n*=48). One subject with chronic HF and two healthy subjects did not provide a blood sample during the study, resulting in a total of 133 data points. PERMANOVA was employed to investigate if there were global cohort-specific cytokine shifts. To identify individual cytokines that were differentially abundant in chronic HF versus healthy subjects, multivariate LMMs accounting for age, sex, race, and BMI as fixed effects and subject as a random effect were employed.

#### Chronic HF phenotype, severity, and clinical change

A multivariate LMM was employed that accounted for subject age, sex, race (white versus other), BMI, and diet as fixed effects, with subject as a random effect. Each multi-omic feature was modeled as a dependent variable, with different HF phenotype, severity, poor clinical outcome or clinical change variable modeled as independent (outlined under “Metagenomic data”). For the volcano plot of circulating cytokines, metabolites, and lipids associated with a poor clinical outcome, *p*-values were calculated from the multivariate LMM, with log_2_(fold change) calculated for mean values of each feature according to the group assignment (poor clinical outcome versus no clinical outcome).

### Microbiome-host interactions

#### Cytokine-microbiome correlations for chronic HF versus healthy individuals

Pairwise Spearman correlations between the most abundant microbial pathways (*n*=100) and cytokines (*n*=76) were calculated for patients with chronic HF (*n*=58), and hierarchical clustering was performed (at both microbiome pathway and cytokine levels) for these samples. The hierarchical cluster structure was then applied to the pairwise Spearman correlations between microbiome pathways and cytokines within the healthy cohort (*n*=48). This procedure revealed a microbiome pathway cluster with significant cohort-specific correlations with cytokines, comprising five pathways for arginine and ornithine biosynthesis (PWY-7400, ARGSYN-PWY, ARGSYNBSUB-PWY, PWY-5154, and GLUTORN-PWY). The relative abundance of these pathways was aggregated, and the Spearman correlation with TNF-⍺, the chief pro-inflammatory cytokine, was calculated. For visualization purposes, aggregated pathways were log_2_-transformed.

#### Bifidobacterium association with host multi-omic biomarkers

*Bifidobacterium* relative abundance was modeled as an independent variable in multivariate LMMs, which accounted for the aforementioned demographics and dietary variables, and subject as a random effect. Similar analyses were performed for *Bifidobacterium* species.

### Machine learning using multi-omic data for classification between chronic HF and healthy state

This analysis excluded features that were only available for one of the cohorts (e.g., NT-proBNP was only available for patients with chronic HF) or where batch effects confounded HF and healthy cohorts (e.g., metabolomics and lipidomics features, which were unavailable for healthy subjects). This filtering left four data types for the analysis: laboratory tests (*n*=53), cytokines (*n*=76), microbial taxa (*n*=553), and pathways (*n*=306). Each of 988 variables was first standardized (zero centered and scaled) among 133 subjects (85 heart failure visits for 58 unique HF patients and 48 healthy subjects), after which 80% of the samples were randomly selected from stratified groups as the training set. The remaining 20% of samples served as the holdout set. With healthy subjects labeled as 0 and patients with chronic HF labeled as 1, Lasso and elastic-net models were fitted for regression on standardized multi-omic variables using coordinate descent as implemented in the R ‘glmnet’ and ‘caret’ packages. To derive feature importance, the coefficients from the model trained with all four-omics types together were used.

### Multiple hypothesis testing

The Benjamini-Hochberg (BH) method was used to correct for multiple hypotheses testing^88^, with adjusted *p*-values (p_adj_)<0.3 considered statistically significant. Analyses for which we did not perform BH adjustment included PERMANOVA, cytokine-microbiome correlation (due to its exploratory nature), and exometabolomics analyses.

### Code

All code was written in R (v. 4.2.3) and is available with the manuscript.

## ACKNOWLEDGMENTS

The authors thank all study subjects for their participation and commitment, as well as the Snyder, Huang and Sonnenburg labs for helpful discussions. Our work was supported by NIH Awards F32HL143916 and K08HL164881 (to P.M.), and K08ES028825 (to S.M.S-F.R.), American Heart Association Career Development Award 856341 (to P.M.), a James S. McDonnell Postdoctoral Fellowship (to H.S.), NIH RM1 Award GM135102 (to K.C.H.), and NIH 5U54HG012723 (to M.P.S.). J.L.S. and K.C.H. are Chan Zuckerberg Biohub Investigators.

## AUTHOR CONTRIBUTIONS

P.M. and M.P.S. conceived the study and designed the experiments. P.M., K.C., and M.A. performed experimental bench work. P.M. conducted host data acquisition and processing. P.M., H.S., W.Z., and M.A. conducted microbiome data acquisition and processing. P.M., and H.S. analyzed the data. P.M. performed integrative data analyses, managed sample inventory, clinical study visits and Stanford IRB, performed data coordination and submission, and managed daily project activities. Additional specific contributions include machine learning analyses (W.Z.), metabolomics data processing and analysis (N.B.), dietary data processing and analysis (H.P.), clinical data processing for the healthy cohort (S.M.S.-F.R.). J.L.S., K.S., W.H.W.T, M.B.F., P.A.H., and K.K.K. advised on the study design and interpretation of results and assisted with obtaining funding. P.M. and M.P.S. obtained funding. P.M. and H.S. wrote the manuscript with critical edits by M.P.S. and K.C.H. and input from all authors.

