## Supplementary Materials for "Gut microbiome shifts in chronic systolic heart failure are associated with disease severity and clinical improvement"

### SUPPLEMENTARY FIGURES

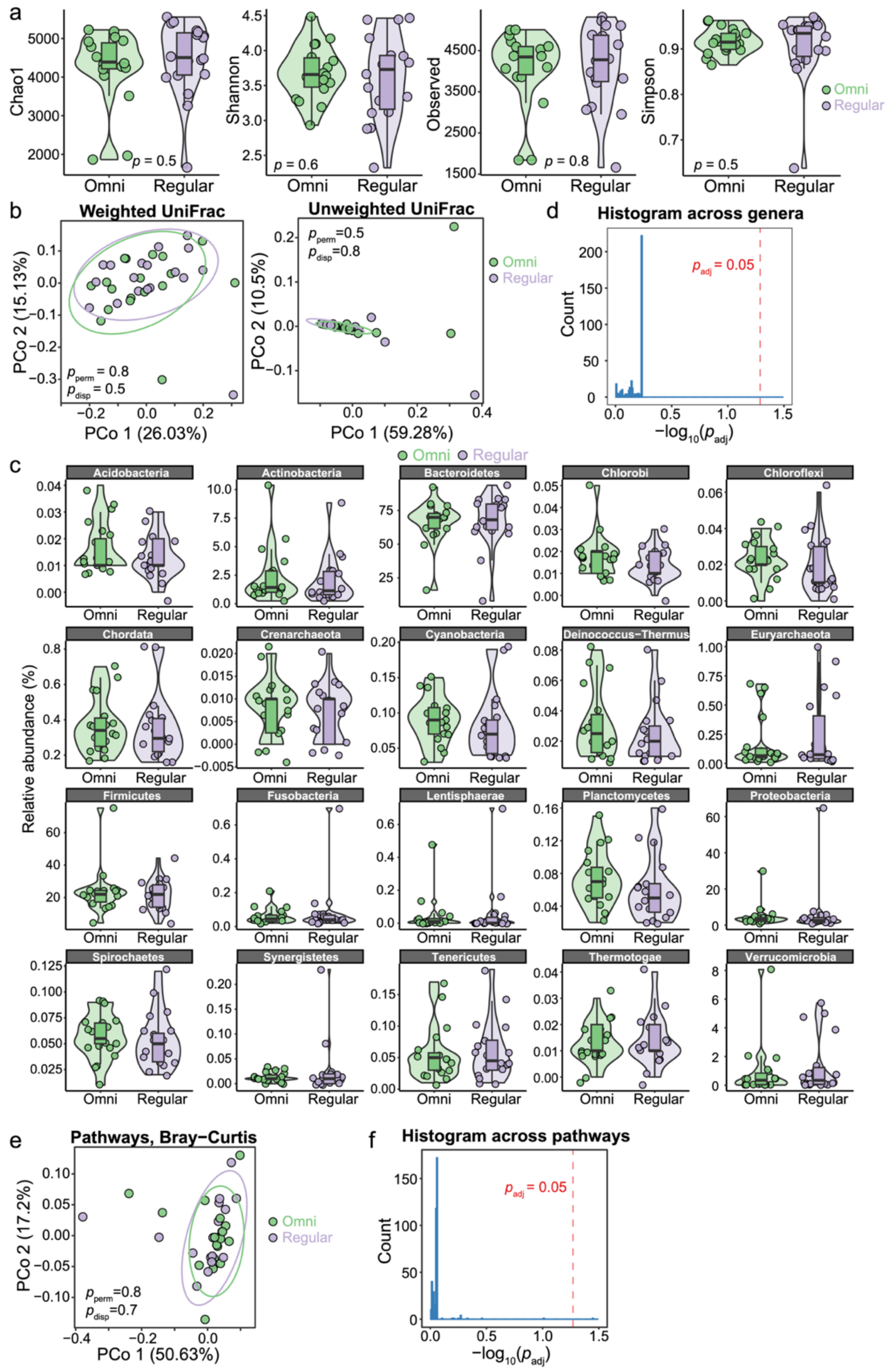

**Figure S1: No significant batch effects were observed between fresh frozen stool samples and samples preserved using OMNIgene-GUT collection kit.** For 18 participants, we collected stool samples using simultaneous direct freezing and OMNIgene-GUT collection kit. **a-d**, The paired samples exhibited no significant differences in alpha (**a**) and beta (**b**) diversities at the taxonomic level. We further used a univariate linear model to evaluate differences in at phylum (**c**) and genus (**d**) level and did not observe any significant differences in either. **e,** Similarly, we did not observe any differences in pathway beta diversity. **f**, When using a univariate linear model to evaluate potential bias from sampling methods in pathway abundances, we found that the teichoic acid (poly-glycerol) biosynthesis pathway (TEICHOICACID-PWY; $p_{adj}$=0.04) and L-lysine biosynthesis II pathway (PWY-2941; $p_{adj}$=0.098) were the only two pathways affected by sampling methods. For downstream analyses, we excluded these two pathways.

$p_{adj}$: *p*-values after FDR correction using the Benjamini-Hochberg method; $p_{perm}$: *p*-values from Permutation Based Analyses of Variance (PERMANOVA); $p_{disp}$: *p*-values from beta dispersion analyses.

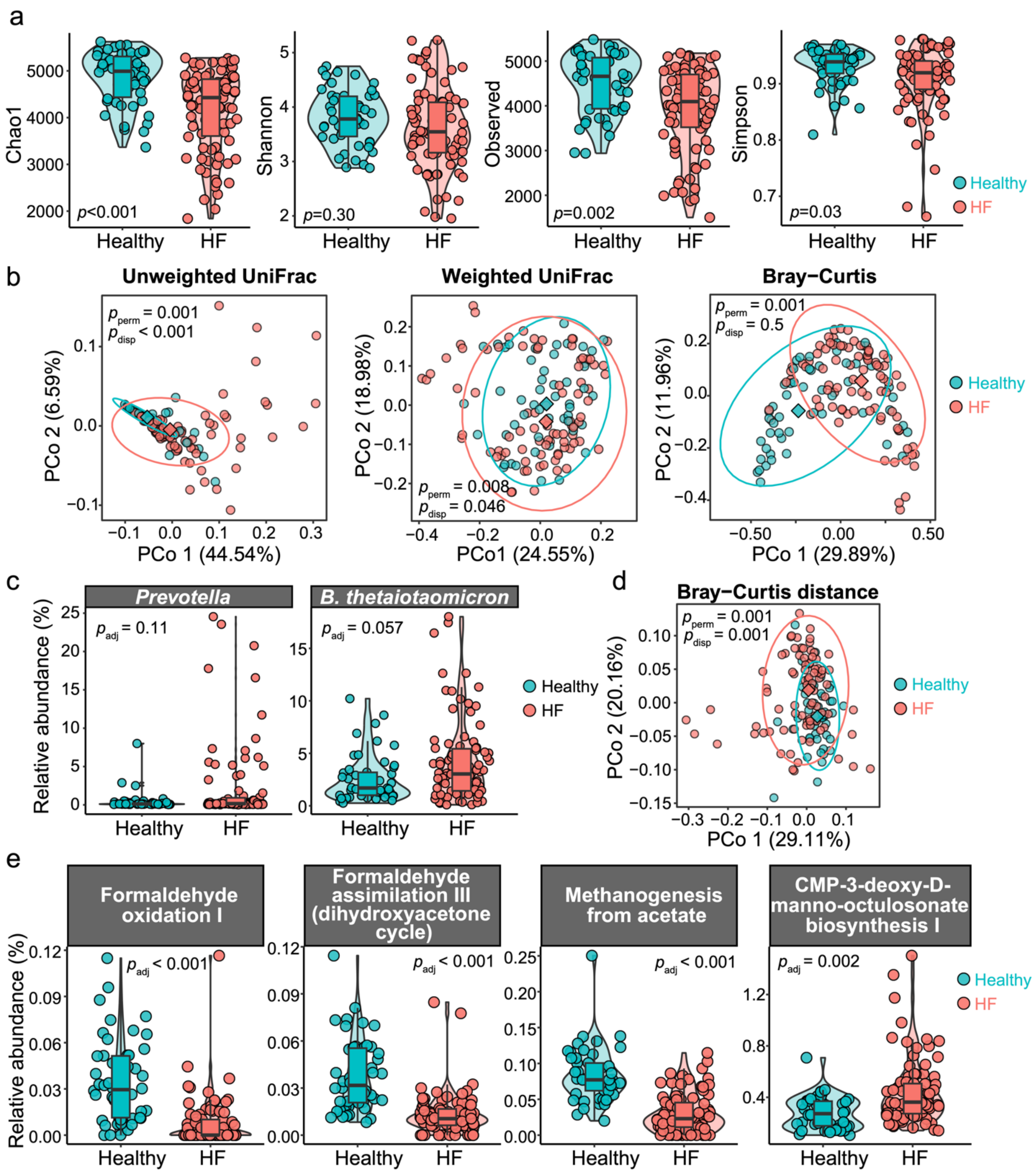


**Figure S2: Chronic heart failure is associated with distinct taxonomic and functional gut microbiome changes. a**, HF subjects had overall lower alpha diversity compared to the healthy cohort, as measured by Chao1 index, number of observed species, and Simpson index. There was no significant difference in Shannon diversity between the two groups. **b**, beta diversity of healthy and HF subjects in taxonomic compositions. The two groups had significantly different compositions measured by unweighted and weighted UniFrac distance, as well as Bray-Curtis distance. **c**, Representative taxonomic groups (*Prevotella* and *Bacteroides thetaiotaomicron*) were differentially abundant in healthy and HF subjects. **d**, Beta diversity in pathways. Healthy and HF subjects had different pathway compositions. **e**, Representative pathways that were enriched or depleted in HF subjects.

$p_{adj}$: *p*-values after FDR correction using the Benjamini-Hochberg method; $p_{perm}$: *p*-values from Permutation Based Analyses of Variance (PERMANOVA); $p_{disp}$: *p*-values from beta dispersion analyses.

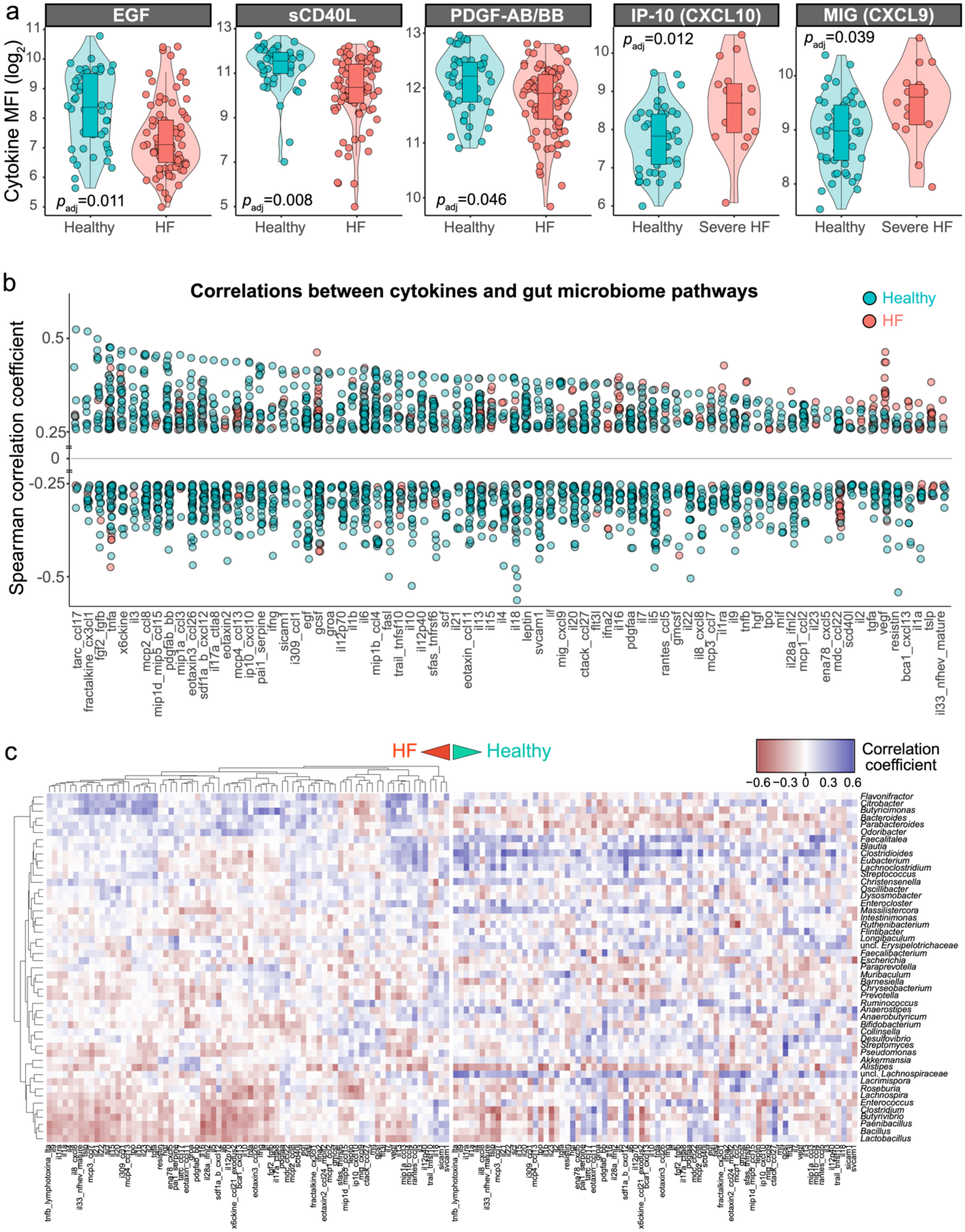
a
EGF
sCD40L
PDGF-AB/BB
IP-10 (CXCL10)
MIG (CXCL9)
Cytokine MFI (log2)
p_adj=0.011
p_adj=0.008
p_adj=0.046
p_adj=0.012
p_adj=0.039
Healthy
HF
Severe HF
b
Correlations between cytokines and gut microbiome pathways
Healthy
HF
Spearman correlation coefficient
c
HF
Healthy
Correlation coefficient
−0.6 −0.3 0 0.3 0.6
Flavonifractor
Citrobacter
Butyricimonas
Bacteroides
Parabacteroides
Odoribacter
Faecalitalea
Blautia
Clostridioides
Eubacterium
Lachnoclostridium
Streptococcus
Christensenella
Oscillibacter
Dysosmobacter
Enterocloster
Massilistercora
Intestinimonas
Ruthenibacterium
Flintibacter
Longibaculum
uncl. Erysipelotrichaceae
Faecalibacterium
Escherichia
Paraprevotella
Muribaculum
Barnesiella
Chryseobacterium
Prevotella
Ruminococcus
Anaerostipes
Anaerobutyricum
Bifidobacterium
Collinsella
Desulfovibrio
Streptomyces
Pseudomonas
Akkermansia
Alistipes
uncl. Lachnospiraceae
Lacrimispora
Roseburia
Lachnospira
Enterococcus
Clostridium
Butyrivibrio
Paenibacillus
Bacillus
Lactobacillus

**Figure S3: Individual cytokines are differentially abundant in patients with chronic HF compared to healthy subjects. a**, Baseline differences in the most differentially abundant cytokines between patients with chronic HF (*n*=58) compared to healthy subjects (*n*=48). Additional differences were identified when only the subset of patients with more severe chronic HF (New York Heart Association class III or IV, *n*=14) were compared to healthy subjects. A multivariate linear mixed model accounting for age, sex, race, and body mass index as fixed effects and subject as a random effect were employed. The Benjamini-Hochberg method was used to adjust for multiple hypothesis testing. **b**, Healthy subjects, compared to patients with chronic HF, had stronger and more numerous correlations between cytokines and gut microbiome pathways. Each point represents a pairwise Spearman correlation between a cytokine (*x*-axis; *n*=76) and individual microbial pathway (*n*=306). Only correlations with absolute correlation coefficient >0.25 are shown. **c**, Heatmap of correlations between the top 50 most abundant microbial genera (rows) and cytokines (*n*=76; columns). Pairwise Spearman correlations were calculated, then hierarchical clustering was performed (both on microbiome and cytokine levels) for the samples from the HF cohort (*n*=58, left). The hierarchical cluster structure was then applied to the pairwise Spearman correlations between microbiome genera and cytokines within the healthy cohort (*n*=48, right).

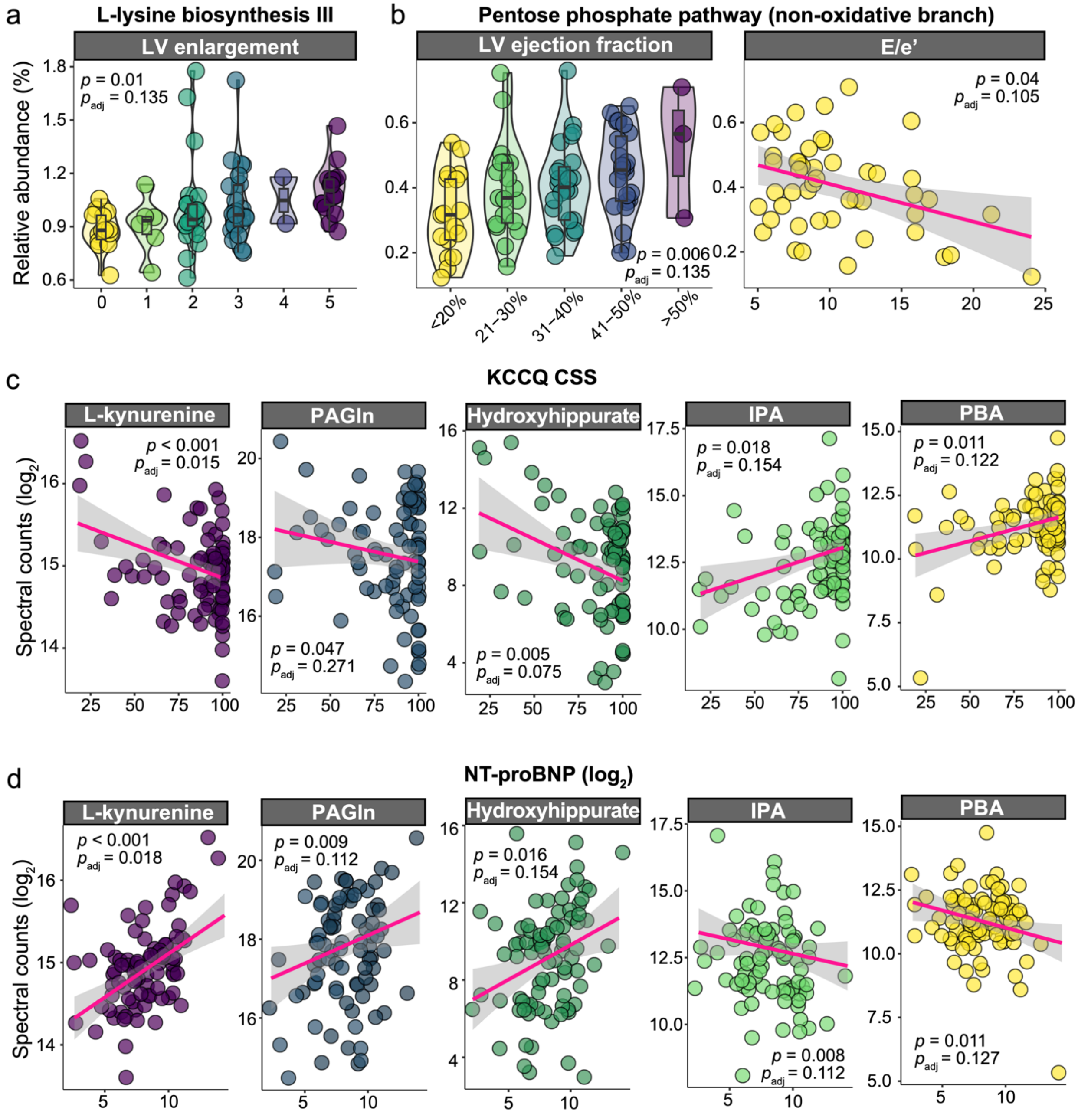


**Figure S4: Gut microbiome functional community features and metabolites are associated with metrics of HF and cardiomyopathy severity. a**, L-lysine biosynthesis is directly associated with the degree of left ventricular (LV) enlargement (0=normal LV size; 5=severe LV enlargement). **b**, Non-oxidative branch of the pentose phosphate pathway is significantly associated with better cardiac systolic (LV ejection fraction) and diastolic function (E/e’). **c**, Multiple circulating gut microbiome-produced metabolites are associated with Kansas City Cardiomyopathy Questionnaire Clinical Summary Score (KCCQ CSS) and N-terminal pro-brain natriuretic peptide (NT-proBNP; $log_2$ transformed). L-kynurenine, phenylacetylglutamine (PAGln), and hydroxyhippurate spectral counts are associated with lower CSS and higher NT-proBNP (indicators of more severe HF), whereas indole-3-propionic acid (IPA) and phenylbutyric acid (PBA) were associated with higher CSS and lower NT-proBNP (consistent with less severe HF). (**a-c**) In a linear mixed model, which accounted for age, sex, race (white versus other), body mass index and dietary patterns as fixed effects, and subject as a random effect, each gut microbiome feature was modeled as a dependent variable, with each of the metrics of HF severity modeled individually as an independent variable. The Benjamini-Hochberg method was used to adjust for multiple hypothesis testing. Abbreviations: LV, left ventricular; E/e', early mitral inflow velocity (E) / early diastolic mitral annular velocity (e'); KCCQ OSS, Kansas City Cardiomyopathy Questionnaire Clinical Summary Score; NT-proBNP, N-terminal pro-brain natriuretic peptide; PAGln, phenylacetylglutamine; IPA, indole-3-propionic acid; PBA, phenylbutyric acid.

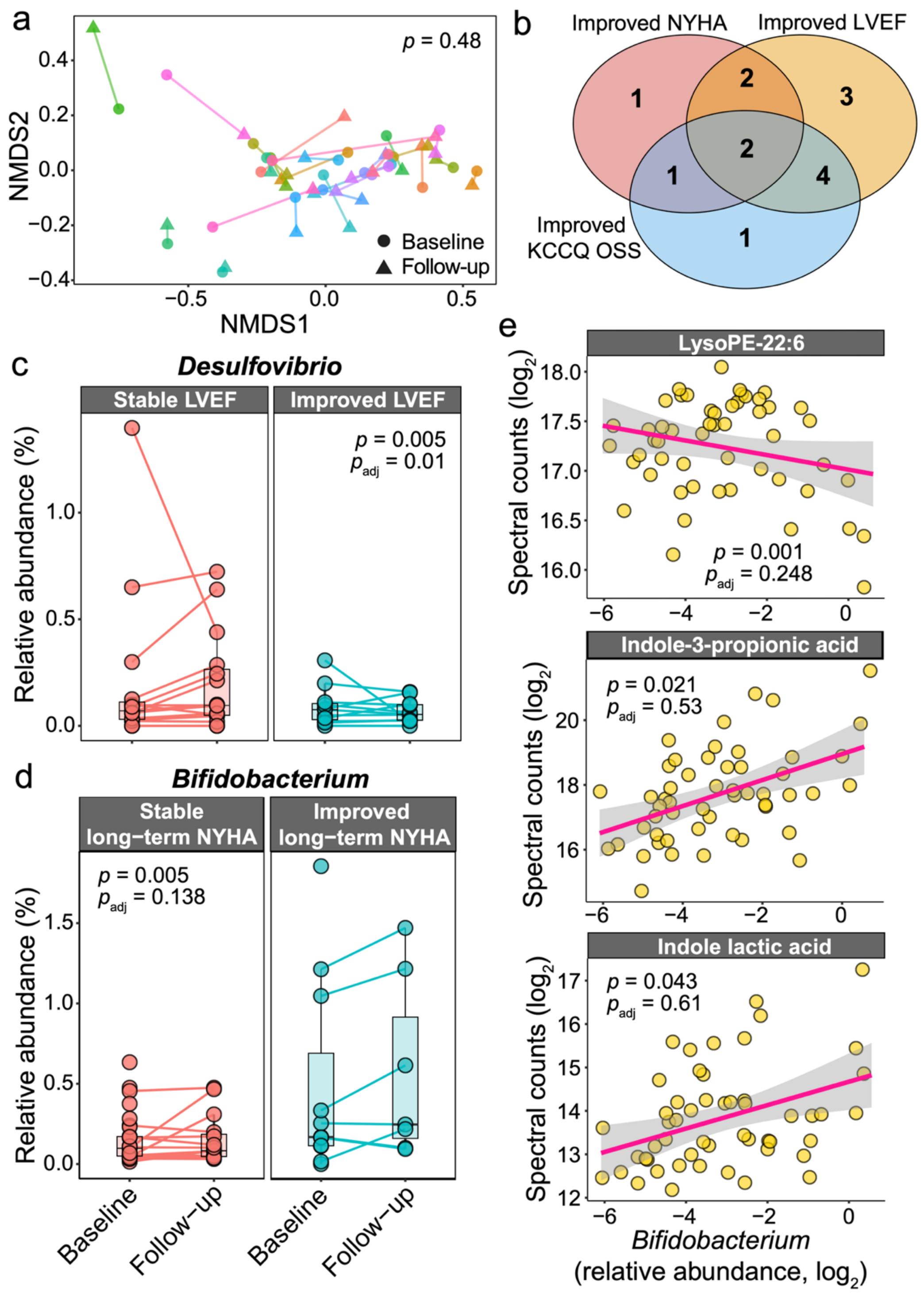


**Figure S5: Bifidobacterium is associated with HF long-term clinical improvement.**
**a**, Gut microbiome beta diversity was not significantly different between baseline and follow-up visits (PERMANOVA $p>0.05$; microbiome pathway comparison shown,

visualized as non-metric multidimensional scaling ordination). Instead, samples from the same subject (different colors) clustered together. **b**, Of the 14 patients who experienced either improvement in the New York Heart Association (NYHA) functional class, left ventricular ejection fraction (LVEF), or Kansas City Cardiomyopathy Questionnaire Overall Summary Score (KCCQ OSS), only 2 patients experienced improvement in all 3 metrics. **c**, *Desulfovibrio* abundance was negatively associated with LVEF improvement over time (*n*=11). **d**, *Bifidobacterium* abundance was associated with long-term NYHA improvement from baseline (*n*=7 of 51 patients with long-term clinical follow-up data available, obtained through the electronic medical record). **e**, *Bifidobacterium* abundance was inversely associated with lysophosphatidylethanolamine (LysoPE-22:6), and nominally directly associated with indole-3-propionate (IPA) and indole lactic acid (ILA) spectral counts. (**c-e**) A linear mixed model accounting for age, sex, race (white versus other), body mass index and dietary patterns as fixed effects, and subject as a random effect was performed, with the Benjamini-Hochberg method used for multiple hypothesis testing. (**c-d**) HF clinical change metric was modeled as an independent variable, with gut microbiome feature modeled as a dependent variable. In **e**, microbiome feature was modeled as an independent variable and metabolites as a dependent variable. Abbreviations: PERMANOVA: permutational multivariate analysis of variance; NYHA, New York Heart Association; LVEF, left ventricular ejection fraction; KCCQ OSS, Kansas City Cardiomyopathy Questionnaire Clinical Summary Score; LysoPE-22:6, lysophosphatidylethanolamine 22:6; IPA, indole-3-propionate; ILA, indole lactic acid.

### SUPPLEMENTARY TABLE CAPTIONS

**Table S1: Clinical variables and baseline comparisons between the heart failure and healthy cohorts.**

**Table S2: Analysis of stool collection method-related batch effects.** Eighteen healthy subjects concurrently provided stool samples via the conventional method and the OMNIgene-GUT collection kit (Methods). These samples were analyzed (*n*=36) and microbiome (phylum, genus, and pathway) signatures were compared.

**Table S3: Chronic heart failure-associated gut microbiome signatures.** Analyses of family, genus, and species taxonomic ranks, metabolic pathways, systemically over- and under-represented microbiome functions, and predicted microbiome-produced metabolites.

**Table S4: Chronic heart failure-associated cytokine signatures.** Between-cohort differences in baseline cytokines as a whole, as well as individual cytokines from all visits (analyzing the entire heart failure (HF) cohort and including only patients with severe HF (New York Heart Association functional class III or IV)).

**Table S5: Associations between gut microbiome features (family, genus and species taxonomic rank, metabolic pathways, and predicted metabolites) and individual metrics of chronic heart failure and cardiomyopathy severity.**

**Table S6: Associations between host circulating multi-omics (cytokines, untargeted metabolites, and targeted lipids) and individual metrics of chronic heart failure and cardiomyopathy severity.**

**Table S7: Heart failure cohort-level between-visit changes in gut microbiome and circulating host multi-omic features.** Analyses evaluating longitudinal (from baseline to follow-up visit) changes in gut microbiome (family, genus, species and pathway level) and host circulating multi-omic features (cytokines, untargeted metabolites, and targeted lipids).

**Table S8: Longitudinal change in heart failure clinical status and association with gut microbiome features.** Analyses examining associations between heart failure clinical improvement (improvement in New York Heart Association functional class, left ventricular ejection fraction, or Kansas City Cardiomyopathy overall summary score) for time interval between baseline and in-person study follow-up visit, and between baseline and long-term follow-up (long-term data extracted from electronic medical record), and gut microbiome features (family, genus, species, and pathway levels).

**Table S9: Associations between the *Bifidobacterium* species and chronic heart failure clinical change over time and individual metrics of heart failure and cardiomyopathy severity.** Four *Bifidobacterium* species were detected in the gut microbiome of our cohort (*B. longum*, *B. breve*, *B. pseudocatenulatum*, and *B. adolescentis*).

**Table S10: *Bifidobacterium* associations with host multi-omic features.** Analyses examining the associations between *Bifidobacterium* abundance in the gut microbiome and circulating host multi-omics (cytokines, untargeted metabolites, and targeted lipids).
